# Host control and regulation of human anellovirus persistence

**DOI:** 10.64898/2026.11.05.764726

**Authors:** Jacob C. Gutierrez, Thahmina A. Ali, C. Cole Honeycutt, Jun Hao Neo, Konstantin Helmsauer, Sherry S. Nyeo, Ann Le, Allison J. Vela, Laura C. Kida, Austin A. Varela, Rico C. Ardy, Eli Metzner, Jacob C. Ulirsch, Ryan S. Dhindsa, Petko P. Fiziev, Rayan Chikhi, Thomas M. Norman, Kushal K. Dey, Caleb A. Lareau

## Abstract

Anelloviruses are the most abundant and diverse eukaryotic viruses known to infect humans and can persist throughout our lifetimes. Despite their ubiquity, the host factors governing their physiologic distribution and abundance remain poorly understood. Here, we construct a viral pangenome of 2,023 strains to quantify anellovirus DNA in >1.1 million participants spanning three genetic biobanks. Genome-wide association studies identify 121 loci linked to viral persistence, predominantly via immune regulation by NF-κB signaling. Further, we associate variable anellovirus detection with conditions characterized by immune dysfunction, including lymphoid malignancies. By screening 3.2 petabases of human single-cell sequencing data, we identify myeloid progenitors in the bone marrow as a prominent site of anellovirus transcription in vivo. Finally, Perturb-seq of anellovirus proteins reveals that anellovirus ORF2, particularly from Alphatorquevirus, attenuates TNF-α-induced NF-κB signaling, connecting host genetic variation, cellular context, and viral immunomodulation to characterize the persistence of the eukaryotic human virome’s most abundant component.

## INTRODUCTION

Human physiology is shaped by a diverse community of ∼10 trillion virus particles that persist within the body but cause no overt acute symptoms, collectively known as the human virome^1^. The most abundant and diverse members annotated in the human virome are species of the *Anelloviridae* family, which comprises small non-enveloped viruses with circular, negative-sense single-stranded DNA genomes of 1.7-3.9 kilobases (kb).^2^ Anelloviruses have been detected in nearly every tissue compartment^3^, and profiling of isolated cell populations has indicated hematopoietic tropism^4^. Our working understanding of these viral species has categorized them into three principal genera denoted *Alphatorquevirus* (or torque teno virus [TTV], 3.9 kb), *Betatorquevirus* (torque teno mini virus [TTMV], 2.9 kb), and *Gammatorquevirus* (torque teno midi virus [TTMDV], 3.2 kb; **Fig. 1A**). First identified as a candidate agent of post-transfusion hepatitis of unknown etiology^5^, anelloviruses were soon identified in healthy donors worldwide, as nearly all viral DNA profiled from the plasma of healthy individuals originates from anelloviruses^6^, and subsequent analyses have failed to assign a causal role in any common disease in humans^7,8^. Instead, our current understanding is that anelloviruses initially infect humans in infancy, colonize nearly every sampled body compartment, and are carried for life without apparent symptoms in most individuals^9,10^. How these viral species persist in the human body and whether their regulation reflects the potential for chronic pathology remain poorly understood.

**Figure 1.**
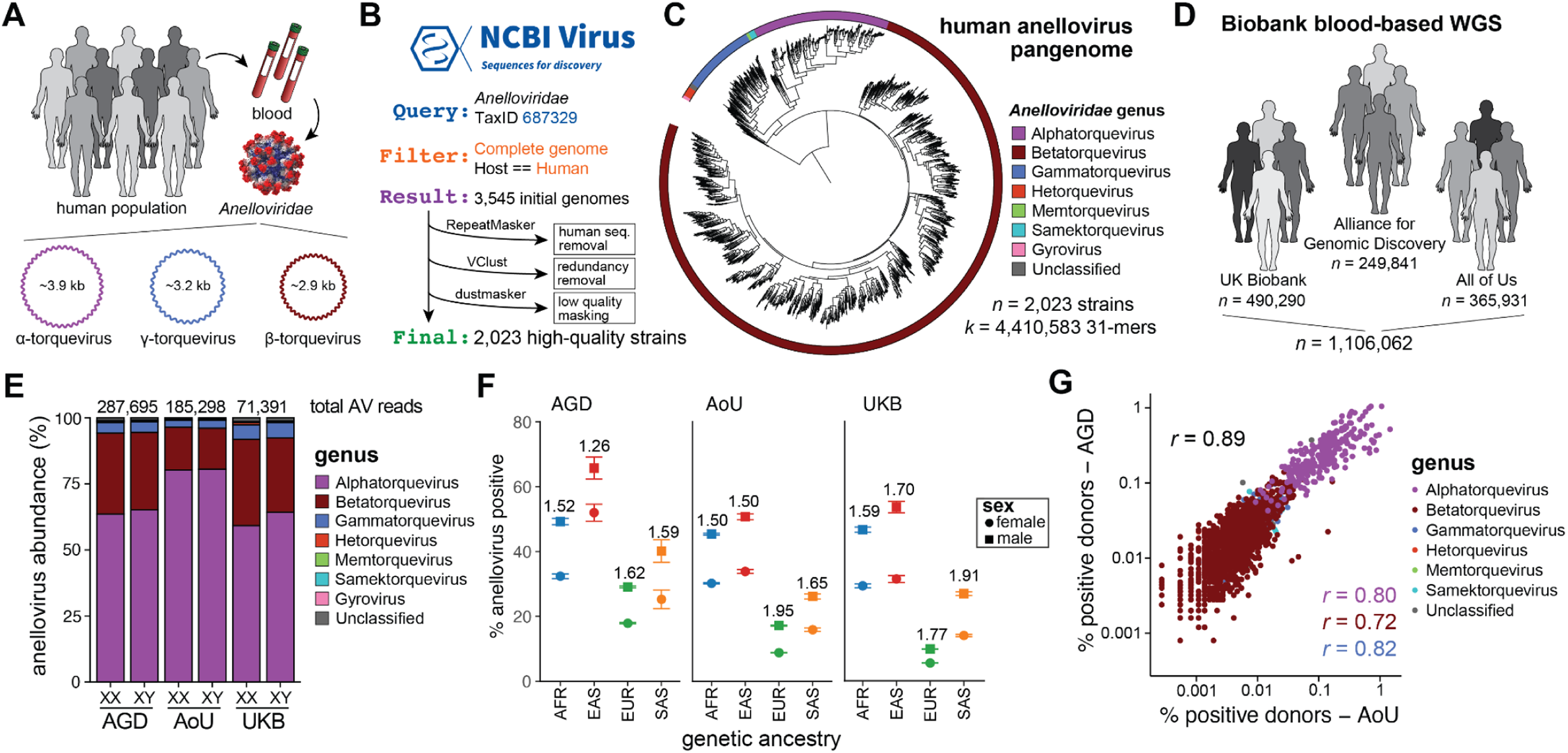
Study design and quantification of anelloviruses across population biobanks. A, Schematic of study design and definition of key aspects of anellovirus biology. **B**, Workflow to generate human anellovirus pangenome. **C**, Visualization of anellovirus strains in the human anellovirus pangenome, colored by the inferred genus of each strain. **D,** Overview of samples used for anellovirus quantification across three distinct genetic biobanks, collectively comprising over 1.1 million individuals. **E,** Summary of total anellovirus (AV) reads in each biobank, stratified by sex and genus. The three cohorts included the Alliance for Genomic Discovery (AGD), All of Us (AoU), and the UK Biobank (UKB). **F,** Prevalence of anellovirus detection stratified by biobank, sex, and ancestry. Values are the fold increases of male vs. female donors. **G,** Corroboration of strain-level quantification between the AoU and AGD biobanks. Values show Spearman correlation overall (black) and for each major genus (colors).

While there have been preliminary^11,12^ but no conclusive reports of anellovirus infection underlying common disease, emerging association studies indicate that they may directly drive rare diseases or serve as a biomarker of heterogeneous human conditions. The most direct evidence for anellovirus pathogenesis comes from dozens of documented cases of acute promyelocytic leukemia (APL)^13–15^. In each of these cases, a part of the viral genome has been shown to integrate into the *RARA* locus and create an oncogenic viral:host fusion transcript that manifests as leukemia with poor overall outcomes^14^. Viral integration appears to be an extreme case, however, as most viral DNA persists as episomes within a poorly defined cellular reservoir^3,16^. While most healthy individuals have detectable but low levels of anellovirus DNA in peripheral blood that persist for decades^17^, these levels rise dramatically after solid-organ and hematopoietic stem cell transplantation (HSCT), reaching peaks near 10^9^ genome copies per milliliter months after transplantation^4^, secondary to immunosuppression. Because high anellovirus titers are routinely observed in these transplant settings, recent work has evaluated whether anellovirus titers could serve as a marker of immune reconstitution in transplant recipients^18,19^. Additionally, recent studies have associated anellovirus abundance and reactivation with various autoimmune diseases, including systemic lupus erythematosus and rheumatoid arthritis^12^, as well as the acute severity and chronic sequelae of COVID-19^11^. These associations motivate further study of anelloviruses, including efforts to assess their prognostic utility^20^ or potential role in pathogenesis. Alternatively, the capacity for anelloviruses to establish long-term persistence without overt disease has motivated the development of anellovirus-based gene-delivery vectors^21^. Understanding the host-virus interactions that sustain anellovirus persistence could clarify their clinical significance and inform their potential therapeutic use.

Recently, we^22^ and others^23–25^ have shown that variable viral persistence in peripheral blood can be determined by reanalyzing unmapped reads from whole-genome sequencing (WGS) data generated in human biobanks. With over a million individuals now profiled by WGS in widely studied cohorts such as the UK Biobank (UKB), All of Us (AoU), and Alliance for Genomic Discovery (AGD) biobanks, coarse quantifications from unmapped reads can power association studies to determine the genetic basis of viral persistence. This framework was demonstrated in the study of Epstein-Barr Virus (EBV)^22,24,25^, a human herpesvirus that establishes latency in B cells in peripheral blood and whose variable persistence could be explained by dozens of genome-wide loci, primarily at the major histocompatibility complex (MHC) locus.^22^ This framework further confirmed cases where viral persistence was linked with chronic diseases, including autoimmunity, cancer, and neurological conditions whose pathology has been linked to EBV^22,24,25^. Leveraging population-scale variation to infer regulatory factors underlying viral persistence is opportune for virome species such as anelloviruses, as experimental systems that represent the breadth of human anellovirus diversity remain limited^8,16^.

Here, we assemble a comprehensive human anellovirus pangenome from >2,000 high-quality viral strains, comprising 4.4 million 31-mers, to resolve viral abundance and strain composition from WGS data of more than 1.1 million participants profiled in three population-scale genetic biobanks. By quantifying strain-resolved anellovirus abundances, we identify reproducible features of viral persistence across all three cohorts, including a strong polygenic architecture that underlies adaptive immune regulation of persistence mediated by over 120 conditionally independent genetic loci. In addition to a strong heritable phenotype, detection of anelloviruses was associated with various complex traits likely reflecting a shared axis of immune dysfunction. Our comprehensive anellovirus reference further recovers evidence of anellovirus expression from existing single-cell RNA-seq (scRNA-seq) libraries, thereby localizing anellovirus transcription to myeloid progenitors in the bone marrow compartment. Finally, transcriptomic profiling of myeloid cells expressing viral open reading frames (ORFs) revealed an attenuated response to TNF-α-mediated NF-κB signaling, implicating anellovirus ORF2 proteins as a basis in the suppressed anti-viral responses, particularly among the *Alphatorquevirus* genus. Together, our work characterizes host factors regulating the most abundant species persisting in the human virome.

## RESULTS

### Comprehensive quantification of anellovirus across biobanks

We reasoned that studying variation in anellovirus persistence should be particularly feasible using genetic biobanks that use peripheral blood as input material, given the widespread abundance of anelloviruses in WGS data^26^, evidence of hematopoietic tropism^27^, and persistence as DNA^28^. Specifically, we sought to quantify from peripheral blood a species-level estimate of anellovirus strains, including from the principal genera of alpha-, beta-, and gamma-torqueviruses (**Fig. 1A**). In contrast to EBV, which can be adequately represented by a single genome that captures most circulating variation in the human population^22^, we hypothesized that the extensive heterogeneity of anelloviruses could not be adequately captured by any individual or small set of reference sequences for genome-wide association studies.

We queried NCBI Virus for full genomes derived from a *Homo sapiens* host with taxonomy IDs consistent with species in the *Anelloviridae* family, resulting in 3,545 entries that represented the breadth of annotated anelloviruses discovered in human sequencing data (**Fig. 1B; Methods**). Upon closer inspection, we observed that several sequences were either highly similar or had low information content, which could confound interpretation (as we characterized with the EBV genome^22^). To mitigate these issues, we first removed likely human-derived contaminating sequences with RepeatMasker^29^, clustered the contigs with Vclust^30^, and hard-masked low-complexity sequences with dustmasker^31^ (**Methods**). This workflow resulted in a consensus set of 2,023 high-quality strains, comprising 4,410,583 unique 31-mers (**Fig. S1A**). Notably, this represents a 40-fold increase over prior efforts to quantify anelloviruses in biobank-scale cohorts that have selected either 5 (Kamitaki^25^) or 38 (Qiu^23^) sequences as representative of anellovirus diversity.

Next, for each strain, we inferred the genus of each species by extracting the ORF1 sequence and further classified each viral strain using hierarchical clustering (**Fig. 1C; Methods**). A median of 4.7% of each viral strain was masked to remove low-quality sequences in the assembly, including AT-rich regions in the anellovirus genome (**Fig. S1B,C**). This collection of viral sequences was indexed using kallisto^32^ for rapid assignment of large sequencing cohorts, consistent with our past work using k-mer classification for efficient unmapped read processing of herpesviruses (**Methods**)^22,33^. To verify both the sensitivity (i.e., true anellovirus detection) and specificity (i.e., minimal false-positive anellovirus detection) of our reference, we analyzed a set of 1,011 blood-based WGS from the Genotype-Tissue Expression (GTEx) consortium and 2,504 genetically diverse genomes from immortalized lymphoblastic cell lines (LCLs) profiled as part of the 1000 Genomes (1000G) Project (**Fig. S1D**). Because peripheral blood should contain anelloviruses in some individuals, whereas LCLs should be largely negative (aside from potential exposure during cell culture), these datasets served as positive and negative controls for anellovirus detection to assess candidate references for anellovirus read classification (**Methods**). Analyzing the 1000G LCL-derived WGS data assigned 746 and 7,493 reads to the Qiu^23^ and unmasked references, reflecting likely errant assignments, whereas the masked and Kamitaki^25^ references showed minimal anellovirus assignment as expected (2 and 0, respectively; **Fig. S1E**). Next, we tested all four references for detection in blood-derived WGS, where anellovirus DNA is expected, and observed a 36.6% relative increase in the fraction of donors assigned at least one anellovirus read (22.4% versus 16.4%; Fisher’s exact test, *p* = 0.0058; **Fig. S1F**). Further, the inclusion of the human T2T reference decreased anellovirus detection by only 1%, whereas the two LCL-derived reads were corroborated by NCBI BLASTn, supporting the accuracy of our assignments (**Fig. S1G-I; Methods**). While the Kamitaki^25^ reference performed well with relatively few k-mers, we noted that only one representative strain was annotated for both beta-and gamma-torqueviruses, limiting genus-level analyses (**Fig. S1J**). Together, these analyses demonstrate the advantages of our newly developed reference, which we hereafter define as our ‘human anellovirus pangenome’.

Having assembled our comprehensive reference, we turned to three independent population-level biobanks totalling over 1.1 million individuals who were profiled with blood-based WGS (**Fig. 1D**). Across each analysis platform, we reanalyzed 4.1 trillion unmapped reads for potential DNA derived from anelloviruses, spanning 1.2 petabases of previously discarded sequencing data, and classified a total of 544,384 reads as high-confidence anellovirus DNA reads (**Fig. 1E; Methods**). Across biobanks, we detected the three principal genera in peripheral blood but observed a predominance of *Alphatorqueviruses*, consistent with reports of *Beta-* and *Gammatorquevirus* colonization during infancy, with *Alphatorquevirus* ultimately defining the principal component of the adult anellome^34,35^. As all libraries in these biobanks are paired with extensive donor-level metadata, we could link variable detection of anellovirus DNA to donor attributes. Stratifying donors by age confirmed an age-associated accumulation of alphatorqueviruses, with smaller age-associated changes in beta-and gamma-torqueviruses (**Fig. S1K**).

For population-scale association analyses, we classified participants with at least one high-confidence anellovirus-assigned read as DNA-detection positive (**Methods**). While our phenotype captures detectable viral DNA at an individual time point rather than over a lifetime infection status, complementary reports have characterized anellovirus persistence over years of longitudinal profiling^17,36^. Across the three biobanks, males had 26-95% higher detection prevalence than females within the plotted ancestry groups; participants of East Asian genetic ancestry also had higher detection prevalence than the other displayed ancestry groups (**Fig. 1F**). We note that these phenotypes reflect a composite of lifetime viral exposure, acute and chronic immune state, and technical factors as well as potential host genetics. Because anelloviruses are most extensively described in immunosuppressed populations, we evaluated five well-described immunosuppression indicators, none of which explained more than 3.5% of anellovirus-positive UKB participants (**Fig. S1L; Methods**). This is consistent with recent reports of anellovirus detection in individuals with no discernible immunosuppression^11^. To investigate whether anellovirus detection reflected general viral control (i.e., shared host determinants that modulate the persistence of multiple viruses), we analyzed individuals with previously annotated EBV DNAemia and found a 19-47% higher prevalence of anellovirus detection but at a very modest effect size, suggesting disparate regulatory factors underlying viral persistence between EBV and anelloviruses rather than a uniformly elevated viremic state (**Fig. S1M**). In aggregate, per-strain quantifications across biobanks were highly correlated (Spearman *r* = 0.85-0.89; **Fig. 1G; Fig. S1N**). In total, our optimized human anellovirus pangenome enabled a reproducible workflow that scales to process petabytes of data for thousands of anellovirus strains.

### Genetic determinants of anellovirus persistence

Motivated by recent reports of a strong heritable component of EBV persistence^22,24,25,37^, we sought to understand host factors that could regulate anellovirus detection in blood WGS. We first assessed variable outcomes from our anellovirus quantification and found that simple binarization at any high-confidence, non-zero exposure produced the most independent genome-wide significant associations (**Table S1**). Using this optimized binary threshold, we performed 13 genome-wide association analyses across different cohort-by-ancestry strata before meta-analyzing the populations (**Methods**). At a total sample size of 1,043,413 individuals across all genetic ancestries, we identified 121 conditionally independent loci genome-wide, nearly 50 more loci than analyzing individuals of European ancestry alone (**Fig. 2a; Fig. S2A; Methods**). While individual variants in proximity to the major histocompatibility complex (MHC) locus were strongly associated with anellovirus DNA detection, partitioned heritability analyses among individuals of European ancestry indicated a stronger component in the non-MHC (3.7%) than MHC (0.47%) loci, a noticeable difference from the concentration of heritability at the MHC locus (5.7%) for EBV^22^ (**Fig. S2B**). Compared with previous GWAS of anellovirus persistence, our phenotype had 5.4-fold higher estimated SNP heritability outside the MHC, despite substantial overlap in the underlying biobank samples (**Fig. 2b; Methods;** *Z* test p = 8.5×10^-62^). Further, our 121 genome-wide significant loci revealed dozens of new loci compared to the seven previously reported^25^, reinforcing the value of our human anellovirus pangenome (**Fig. 2B,C; Methods**). Gene set enrichment analyses of genes near genome-wide significant loci demonstrated a clear enrichment for pathways implicated in regulatory functions of the immune system, including two pathways linked to TNF signaling (TNFR2 non-canonical NF-κB pathway, adj. *P* = 1.2×10^-4^; TNFs bind their physiologic receptors; adj. *P* = 6.2×10^-3^) as well as B cell signalling (adj. *P* = 0.015; **Methods**) (**Fig. 2D**). Implicated loci included *REL* (encoding c-Rel, a key NF-κB family transcription factor), the *CXCL9-11* chemokines, and *TNFSF8* (**Fig. 2E**). Other significant loci included *C1S*/*C1R* and *CR1*/*CR2*, consistent with prior reports of increased anellovirus loads following complement blockade with an anti-C1S antibody^38^ (**Fig. S2C**). Globally, shared heritability analysis between EBV persistence^22^ and our genetic association here was not related (LD score genetic correlation *r* =-0.11; *p* = 0.13), suggesting disparate mechanisms of regulating viral persistence.

**Figure 2.**
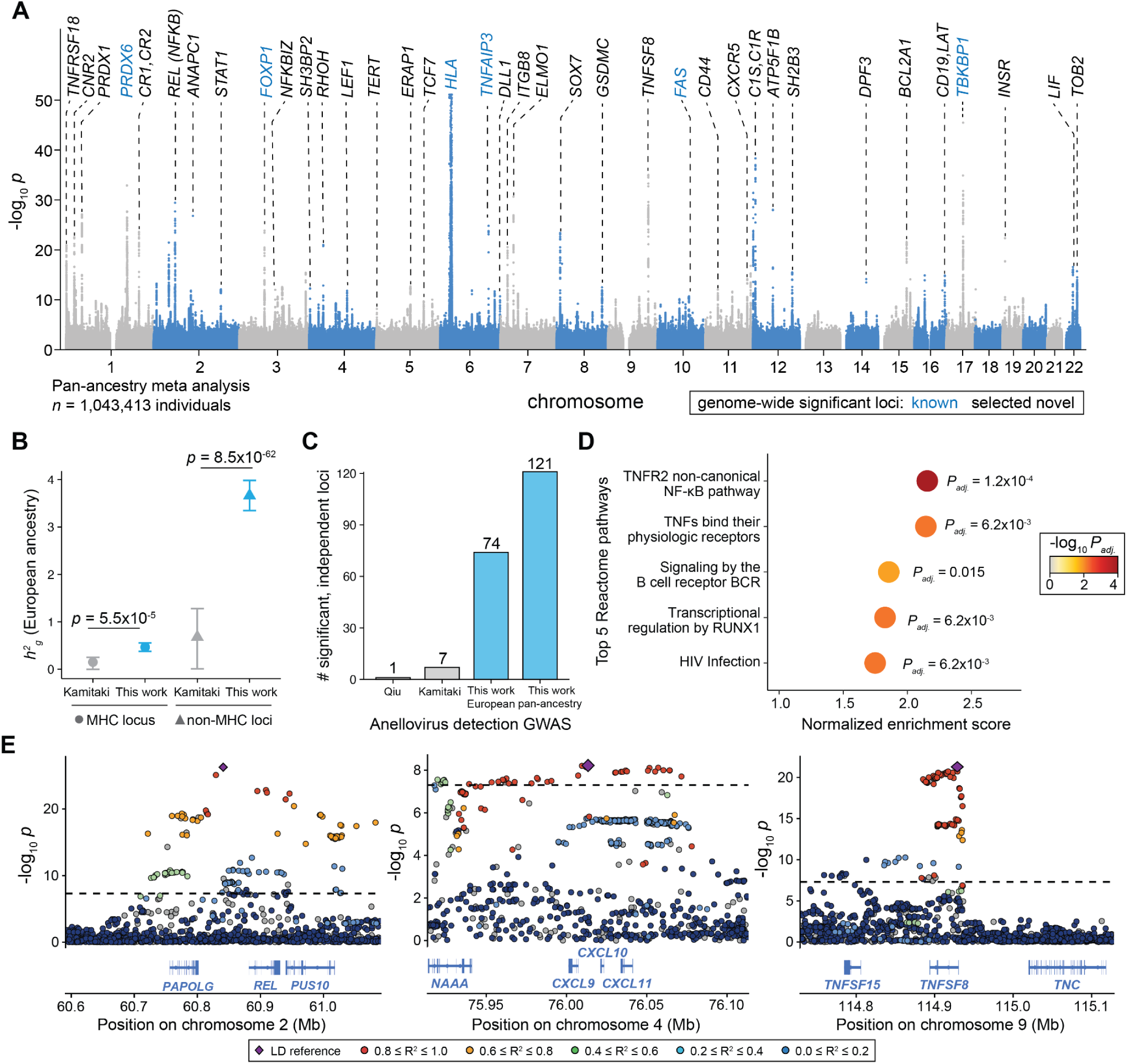
Genome-wide association studies of anellovirus DNA detection. A, Manhattan plot summarizing genome-wide association statistics for anellovirus detection for >1 million individuals, from a meta-analysis across all three biobanks. Genes proximal to selected genome-wide significant associations (*P* < 5 × 10⁻⁸) are annotated. **B,** Comparison of narrow-sense heritability from European ancestry at the MHC locus (left) and outside the MHC locus (right) for the Kamitaki^25^ GWAS and the European GWAS presented here. Statistical test: *Z* test. **C,** Number of conditionally independent, genome-wide significant loci from various anellovirus DNA detection GWASes, including Qiu^23^ and Kamitaki^25^ as well as the European-only and pan-ancestry GWASes presented here. **D,** Top 5 enriched Reactome pathways. Statistical test: MAGMA-based gene set enrichment analysis (GSEA) among Reactome with Benjamini-Hochberg multiple testing correction. **E,** Locus zoom plots for three selected loci, including *REL* (NF-κB), *CXCL9-11*, and *TNFSF8*.

As we previously identified MHC class II variation as the primary driver of persistent EBV infection^22^, we sought to better understand the discordance in the genetic architecture between viruses (**Fig. S2B**). We reasoned that the immense genetic diversity of anelloviruses likely results in disparate peptides that could be presented on specific HLA alleles, whereas EBV has comparably little genetic diversity in the population. To test this, we used the imputed four-digit HLA alleles available for all individuals of European ancestry in the UKB^22^ to train a polygenic risk score (PRS) using the six assigned HLA alleles per donor and tested the accuracy of predicting anellovirus detection in 200,715 EUR donors from AoU (**Methods**). Consistent with the concentration of EBV heritability in the MHC, prediction was more accurate for EBV (AUROC = 0.66) than for the composite anellovirus phenotype (AUROC = 0.57; **Fig. S2D**). To assess whether anellovirus strain diversity contributed to this difference, we repeated the analysis for a well-detected α-TTV strain, OZ258617.1. Prediction improved to an AUROC of 0.69, suggesting that the broad diversity of anelloviruses obscures the strain-specific HLA associations (**Fig. S2D**). Similarly, stratification by HLA-based risk deciles showed stronger enrichment for OZ258617.1 and EBV (OR = 7.19 and 7.57, respectively) than for the composite (OR = 2.45; **Fig. S2E**). HLA-based prediction also extended to other well-represented strains in UKB, and performance correlated with strain-specific sample size (**Fig. S2F**). These findings suggest that the immunodominant epitopes differ across anellovirus strains, and distinct HLA backgrounds similarly enhance or preclude viral persistence. Although HLA-based scores predicted viral presence, they showed no meaningful dose–response relationship with viral read abundance (**Fig. S2G**). Together, these results illustrate how our comprehensive anellovirus pangenome captures both strain-specific HLA associations and broader host regulatory factors outside the MHC that less comprehensive detection approaches missed.

### Multimodal correlates of anellovirus detection

While our genetic analyses supported that detection of any anellovirus was a heritable phenotype controlled by over a hundred host genetic loci, we noted that a subset of donors in each biobank (0.2-2.3%) had considerably higher anellovirus loads (50+ reads; **Fig. 3A**). We hypothesized that such donors may be experiencing an active anellovirus infection where viral transcripts would also be detected. To first verify our human anellovirus pangenome in identifying RNA derived from anelloviruses, we requantified 1,392 RNA-seq libraries that characterized anellovirus reactivation in donors with long COVID^11,39^. Indeed, our k-mer-based quantification closely mirrored the prior anellovirus quantification (*r* = 0.87) but had a 40% boost in classification sensitivity, again reflecting the value of our encompassing viral pangenome (**Fig. S3A**). Next, we quantified the set of 8,980 RNA-seq libraries from the AoU v9 release from donors with paired WGS and assessed viral RNA detection of the same viral strain detected in DNA. Remarkably, we observed a clear dose-response: individuals with higher anellovirus DNA levels had a higher likelihood of transcriptional detection, with up to 98.2% of strains detected at 50+ reads in WGS also detected in RNA-seq (**Fig. 3B**; *p* = 2.7×10^-82^). An analogous comparison using paired WGS and RNA-seq data from GTEx showed a similar trend, where 94.1% of anellovirus strains quantified at 50+ reads in WGS were detectable in RNA-seq data (**Fig. S3B,C**). Conversely, anellovirus strains with only 1 WGS read were detected in only 10-18% of RNA-seq libraries, and these instances made up the majority of anellovirus-positive donors in our biobanks (**Fig. 3A**).

**Figure 3.**
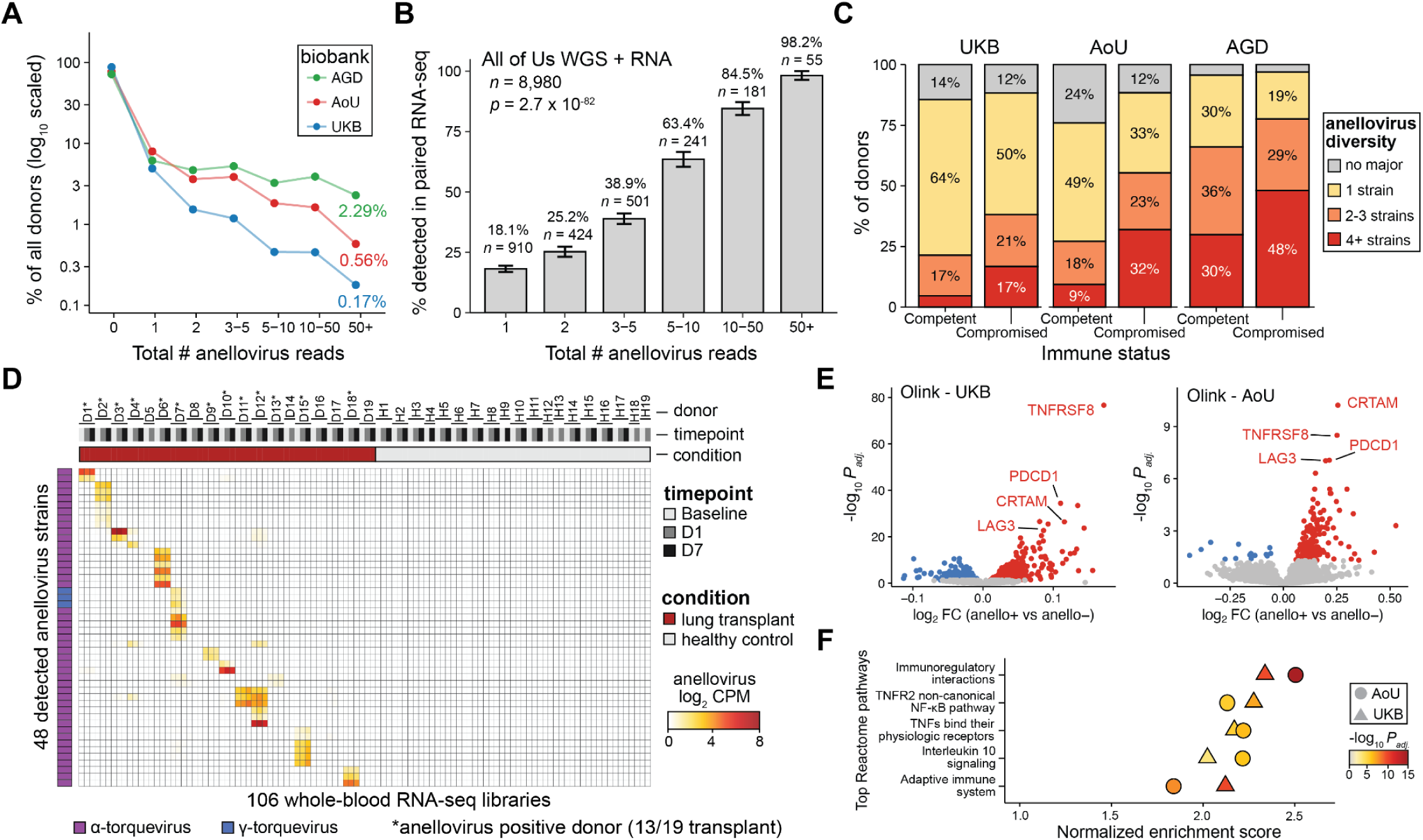
Characterization of anellovirus abundance and diversity. A, Summary of anellovirus read abundances across three biobanks. **B,** Summary of anellovirus transcriptional activity. Binned donor-strain WGS reads were tested for evidence of RNA in matched AoU RNA-seq data. Statistical test: Cochran-Armitage trend test. **C,** Characterization of strain diversity among anellovirus-positive donors, stratified by immunocompromised status in three biobanks. **D,** Characterization of anellovirus expression from RNA-seq libraries of 19 transplant and 19 control donors. Data reanalyzed from prior report.^42^ **E,** Differential plasma protein abundance in the two biobanks, comparing anellovirus-detected and non-detected individuals. **F,** Gene set enrichment analyses of the top 5 Reactome pathways for each differential Olink association set. Statistical test: variance-moderated t-test with Benjamini-Hochberg multiple testing corrections.

We sought to further investigate individuals with 10+ anellovirus reads in each biobank, which comprised 3,172 UKB, 8,711 AoU, and 16,404 AGD donors (**Fig. 3A**). As only 12.8-23.0% of these donors had a clinical annotation of immunosuppression, either from a diagnostic code, transplant, or peripheral blood measurement, we wondered whether the diversity of viral strains, rather than simple abundance, would reflect an outsized effect in immunosuppressed individuals. Leveraging the wealth of strains in our human anellovirus pangenome, we identified whether individuals carried one principal strain or multiple (**Methods**). Across all three cohorts, we observed a 60-240% increase in the proportion harboring 4+ anellovirus strains between immunosuppressed and immunocompetent donors (**Fig. 3C**). This finding suggests that 1-2 predominant strains can persist in immunocompetent individuals, whereas immunosuppression facilitates ‘blooms’ of heterogeneous strains^40,41^. We confirmed that increased anellovirus strain diversity was a feature of immunosuppression after adjusting for relevant covariates, including age, sex, and depth of anellovirus detection (**Fig. S3D; Methods**). Given our findings of anellovirus transcription in AoU and GTEx and elevated strain diversity in immunosuppressed individuals, particularly transplant recipients, we reasoned that these observations would replicate in other cohorts. Indeed, reanalysis of longitudinal RNA-seq profiling of 19 healthy controls and 19 lung transplant recipients showed that a majority (n=13) of transplant recipients had multiple detectable anellovirus strains that persisted over a week of RNA-seq monitoring, whereas no healthy controls had detectable viral RNA^42^ (**Fig. 3D**). Unlike EBV, for which transcriptional activity was rare even at high DNA levels^22^, elevated anellovirus DNA abundance was frequently accompanied by viral RNA detection, often in immunosuppressed individuals with increased strain diversity.

Finally, we sought to characterize whether anellovirus detection could be tied to a consistent proteomic signature in peripheral blood. Plasma proteomics in UKB and AoU yielded concordant signatures in anellovirus-positive donors, including consistent and dose-responsive effects for *TNFRSF8*, *PDCD1*, *CRTAM*, and *LAG3*, markers of T cell dysfunction and chronic antigen exposure (**Fig. 3E; Table S2**). Gene set enrichment converged on five pathways enriched in both biobanks, including immunoregulatory interactions between lymphoid and non-lymphoid cells, TNFR2 non-canonical NF-κB signaling, and TNF receptor binding (**Fig. 3F; Fig. S3E-G**). These findings suggest that anellovirus-positive individuals, particularly those with higher viral loads, exhibit sustained immune activation that may reflect ongoing antigen exposure alongside incomplete viral control. Together, our proteo-genomic characterization corroborates the same axis of adaptive immune control of anelloviruses implicated by our genetic associations, both of which converge on TNF-α-induced NF-κB signaling.

### Clinical associations of anellovirus persistence

Motivated by our consistent genetic and phenotypic correlations, we performed a phenome-wide association study (PheWAS) to map outcomes cataloged via International Classification of Diseases, 10th revision (ICD-10) codes with detected anellovirus DNA as the exposure, analogous to recent work with EBV^22^. Using the UKB non-Finnish European ancestry (NFE) donors as a discovery cohort, we analyzed 12,661 phenotype codes (phecodes) as previously described^43^ (**Methods**). After multiple testing corrections, we identified 58 statistically significant phecodes that collapsed onto 16 unique traits (**Fig. 4A; Methods; Table S3**). Diagnostic codes that remained significant after correction for multiple testing included chronic lymphocytic leukemia, chronic renal failure, and type 2 diabetes. In chronic renal failure and type 2 diabetes, elevated anellovirus detection may reflect impaired antiviral control secondary to chronic immune dysfunction associated with uremia and metabolic dysregulation, respectively. For the 16 significant diagnostic traits, we tested the AoU and AGD biobanks for replication, observing that 13 (81%) replicated in both cohorts (**Fig. 4B; Fig. S4A**, nominal *p* < 0.05; consistent effect direction). Further, we examined whether the genetic architecture uncovered from our association analyses shared significant heritability with these associated traits, which suggested that associations with type 2 diabetes, angina pectoris and anemia, among others, shared a common axis of disease risk with our identified risk axis of anellovirus persistence (**Fig. S4B**). Notably, systemic lupus erythematosus (SLE) showed no clear association, whereas rheumatoid arthritis (RA) showed modest associations in UKB and AoU that did not replicate in AGD (**Fig. 4B**), a departure from the strong associations previously reported in case-control cohorts (previously reported OR = 15.4 for RA and 58.5 for SLE)^12^, which we attribute to the differences in study design, noting biobank studies may mitigate unobserved confounders (**Methods**). Additionally, comprehensive associations of the genetic basis of anellovirus persistence with autoimmune and autoinflammatory traits via cupcake^44^ similarly showed no meaningful association (**Fig. S4C**), suggesting the genetic axis of anellovirus persistence is not strongly shared with the genetic risk of developing these inflammatory diseases.

**Figure 4.**
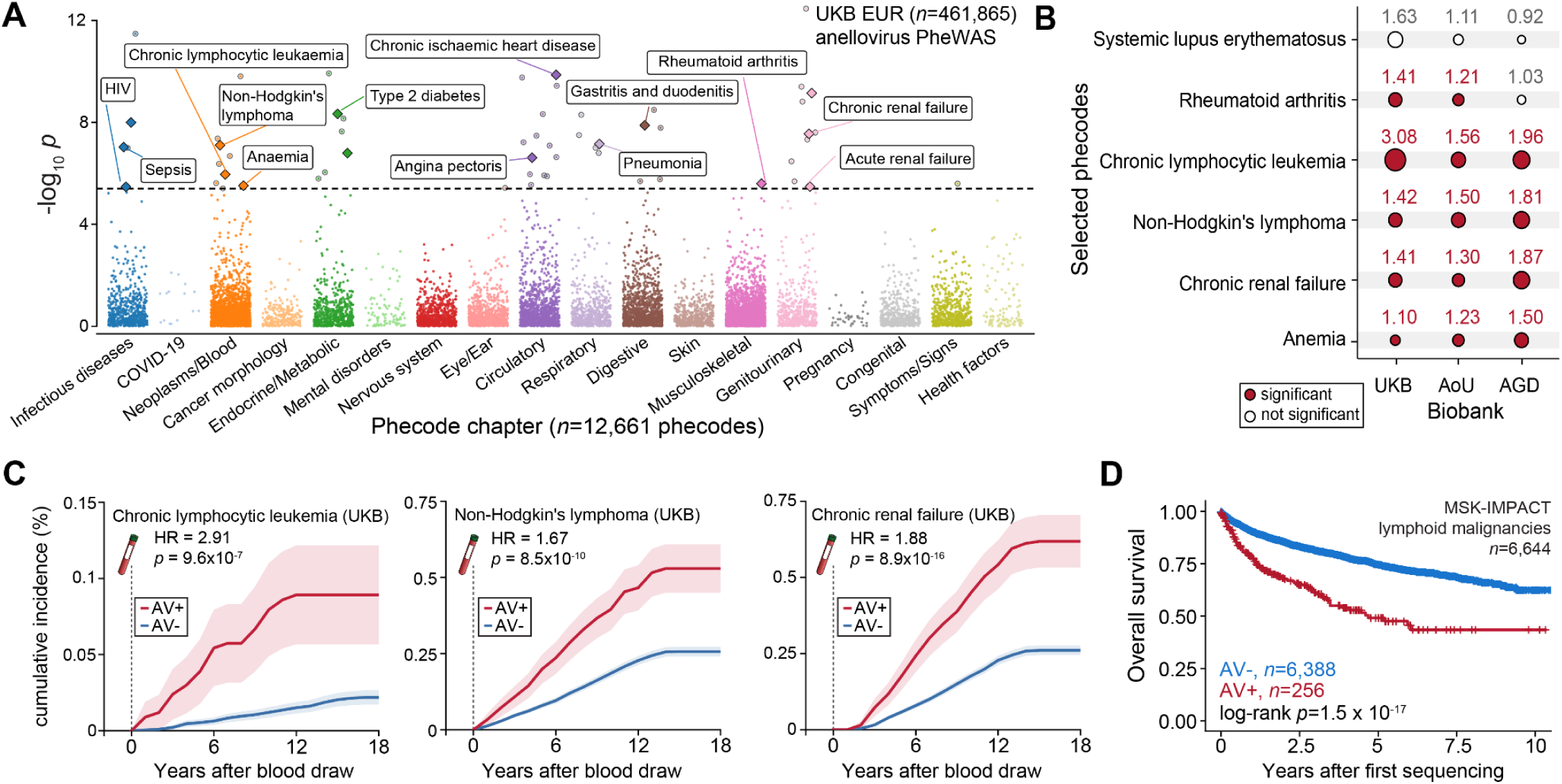
Anellovirus DNA detection associates with complex traits linked to immunosuppressive states. A, Summary of associations between anellovirus DNA detection and binary phenotypic traits in the UKB, with individuals of broadly NFE ancestry. The horizontal dashed line represents the phenome-wide significant threshold (P < 3.95 × 10^-6^). Union phenotypes are plotted to reduce redundancy. Selected traits are highlighted based on biological interest. Statistical test: Wald test from logistic regression model (two-sided). **B,** Comparison of effect sizes and significance for selected traits across the three genetic biobanks. Values represent odds ratios of association, and significance reflects Bonferroni-adjusted associations. **C,** Cumulative accruals of positive diagnostic codes for chronic lymphocytic leukemia (C91.1), non-Hodgkin’s lymphoma (C85.90), and chronic kidney failure (N18). Donors were stratified by anellovirus (AV) detection at the blood draw noted at year 0. Statistical test: Cox proportional hazards model. **D,** Survival outcomes of patients with lymphoid malignancies in MSK IMPACT stratified by detected anellovirus in tumor samples. Statistical test: log-rank test.

While consistent across biobanks, we emphasize that these common diseases are unlikely to be caused by anellovirus infections that are similarly endemic in the population. Instead, we reason that a common risk axis could underlie elevated viral persistence and disease susceptibility, thereby driving the association. We hypothesized that the temporal nature of the UKB data provided a unique opportunity to more deeply characterize our associations and potentially nominate settings where prospective detection of anelloviruses could inform patient stratification. For each phecode, we excluded individuals with a diagnosis before the WGS blood draw and assessed accrual of diagnoses strictly after sample collection (**Methods**). We then used Cox proportional hazards modeling to test whether individuals harboring anellovirus DNA were more likely to be diagnosed with these same chronic conditions. Indeed, for traits such as chronic lymphocytic leukemia (HR=2.91; *p* = 9.6×10^-7^), Non-Hodgkin’s lymphoma (HR=1.67; *p* = 8.5×10^-10^), and chronic renal failure (HR=1.88; p = 8.9×10^-15^), anellovirus DNA stratification was significantly associated with increased disease risk (**Fig. 4C; Fig. S4D**). Notably, this time-to-event model similarly showed no association with SLE (HR=1.04; p = 0.85) and only a modest effect with RA (HR=1.32; *p* = 1.8×10^-9^), further suggesting that elevated anellovirus persistence is not a common driver of autoimmune disease.

Based on associations with lymphoid cancers, we hypothesized that anellovirus detection may further reflect disease severity in patients with these hematologic malignancies. The Memorial Sloan Kettering (MSK)-IMPACT cohort represents one of the largest collections of patients with paired genomic profiling and clinical records^45^. Though DNA is enriched for panels of specific cancer-associated genes, highly prevalent viruses had detectable DNA in these samples^46^. Using our human anellovirus pangenome, we requantified 205,156 sequencing libraries, recovering anellovirus reads in 1.8% of tumors (*n*=117,635) and 1.2% of normal controls (*n*=87,521), with sex-based variation consistent with prior biobank data (**Fig. S4F-H**). We also observed that anellovirus positivity was higher in patients who had already received treatment prior to inclusion in MSK-IMPACT, potentially reflecting disease-and treatment-induced immunodeficiency (**Fig. S4I**). While anellovirus DNA was detectable in some patients across all major cancer types, we observed a clear enrichment in lymphoid malignancies (**Fig. S4J**), consistent with our PheWAS associations (**Fig. 4A**). Further examining lymphoid malignancies, we stratified patients based on their detected anellovirus DNA status. In line with immunodeficiency as a main driver of lymphoma morbidity and mortality, we observed that individuals harboring anellovirus DNA had worse overall survival outcomes compared to individuals with no anellovirus detection (HR = 2.41; log-rank test *p* = 1.5×10^-17^; **Fig. 4D**). This effect was only moderately attenuated (HR = 2.08) after adjusting for pretreatment status, diagnosis, stage, sex, and technical covariates, suggesting that this association does not merely reflect treatment-induced immunodeficiencies or differential anellovirus prevalence across diagnostic groups (**Fig. S4K,L**). We therefore propose that prospective anellovirus DNA detection may help identify patients with impaired immune function and high-risk disease.

### Inference of cell type-resolved anellovirus transcription

Though our analysis of biobank-scale peripheral blood data corroborates the established hematopoietic tropism of anelloviruses, we sought to refine what cell populations harbor anellovirus transcripts. Prior work has implicated a potential anellovirus reservoir in both granulocytes^4^ and T cells^11,47^. We recently demonstrated petabase-scale (10^15^ nucleotides) reanalysis of existing sequencing datasets to uncover previously undescribed viral tropism, including in single-cell RNA-sequencing (scRNA-seq) datasets^33,48^. Given the prevalence of anelloviruses, we reasoned that published scRNA-seq libraries have likely barcoded viral transcripts that were unannotated due to a lack of comprehensive reference. To assess cells harboring anellovirus directly, we processed 3.2 petabases of human scRNA-seq data from accessions present in Logan^49^ and mapped all assembled contigs to our human anellovirus pangenome (**Fig. 5A; Methods**). Alignment of Logan contigs recovered anellovirus sequences in 1,666 single-cell accessions. We ranked accessions by a nomination score that weighs total aligned bases against contig length, separating 91 high-scoring accessions from a tail of 1,575 accessions with only sparse alignments (**Fig. 5B; Methods**). Tissue annotation of the anellovirus-associated accessions was dominated by bone marrow (*n* = 58; Fisher’s Exact Test *p* = 2.1 × 10^-33^), followed by peripheral blood (*n* = 19; Fisher’s exact test *p* = 0.58) and all other tissues (*n =* 14), corroborating hematopoietic tropism of anelloviruses and implicating a bone marrow origin (**Fig. S5A; Table S4**).

**Figure 5.**
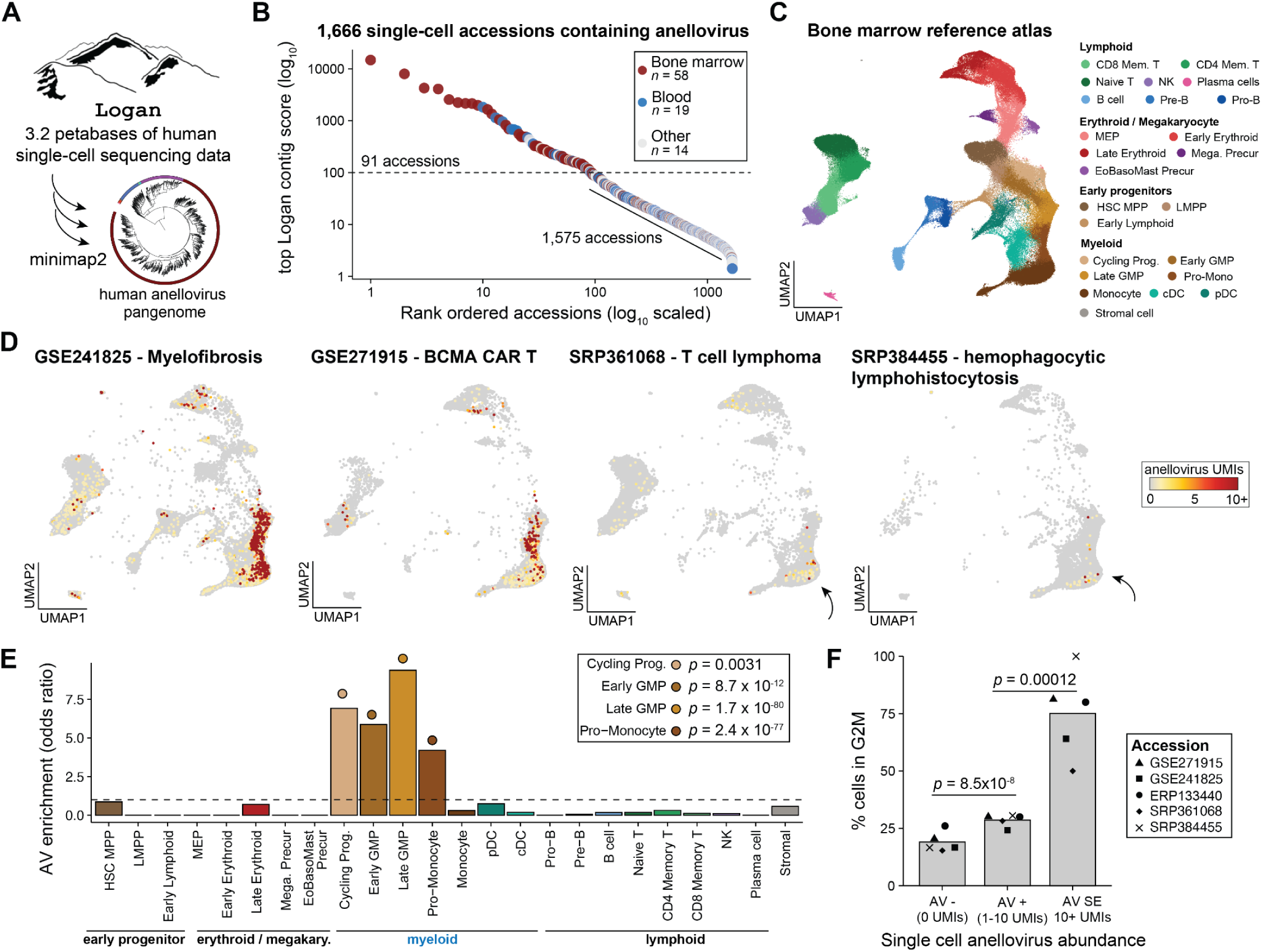
Petabase-scale association study resolves cellular populations underlying anellovirus persistence in the bone marrow. A, Schematic of petabase-scale query using single-cell profiling data and our human anellovirus pangenome reference. **B,** Summary of single-cell profiling accessions with evidence of anellovirus expression. 91 accessions were enriched above a contig score of 100, including 58 from bone marrow and 19 from peripheral blood, as annotated. **C,** Reference embedding of a healthy bone marrow atlas with previously described UMAP coordinates and cell types^50^. **D**, Projection of four bone marrow scRNA-seq datasets showing abundance of anellovirus reads in individual cells. **E,** Summary of specific cell types enriched for anellovirus-positive cells. Statistical test: two-sided Fisher’s exact test. **F,** Percentage of cells in the G2M phase of the cell cycle stratified by abundance of anellovirus (AV) UMIs. Statistical test: Wald test from beta regression.

To identify the specific cells expressing anellovirus in the bone marrow, we retrieved raw sequencing reads for the highest-scoring accessions and processed them along two parallel paths, quantifying anellovirus transcripts against our anellovirus pangenome with kallisto and human transcripts with CellRanger (**Methods**). Libraries derived from bone marrow mononuclear cells (BMMCs) were projected onto the MarrowMap reference atlas to assign harmonized cell type labels and UMAP coordinates (**Fig. 5C**)^50^. We identified five independent studies where BMMCs were profiled spanning myelofibrosis (GSE241825)^51^, BCMA CAR T therapy (GSE271915)^52^, T cell lymphoma (SRP361068)^53^, hemophagocytic lymphohistiocytosis (SRP384455)^54^, and allogeneic HSCT (ERP133440)^55^, where anellovirus-derived UMIs primarily localized to the contiguous trajectory of myeloid differentiation (**Fig. 5D; Fig. S5B**). Anellovirus transcripts in each library were concentrated in a relatively small number of cells, consistent with our characterization of herpesvirus super-expressors (defined by 10+ viral UMIs), and each donor expressed strains that were mostly distinct, consistent with the immense genetic diversity of anelloviruses (**Fig. S5C**). We note the diversity of viruses further minimizes the likelihood of any systematic bias driving the observed expression.

Using the MarrowMap-annotated cell type annotations, we tested each population for enrichment of anellovirus-positive cells (**Fig. 5E; Methods**). Four populations were enriched, all within the myeloid compartment: late granulocyte-monocyte progenitors (GMPs; odds ratio [OR] = 9.2, *p* = 1.7 × 10^-80^), cycling progenitors (OR = 6.9, *p* = 0.0031), early GMPs (OR = 5.9, *p* = 8.7 × 10^-12^), and pro-monocytes (OR = 4.2, *p* = 2.4 × 10^-77^). The enriched cell state expressed *MPO*, *ELANE*, *CES1*, *S100A8*, and *RNASE2*, identifying it as committed granulocytic and monocytic precursors (**Fig. S5D**). Conversely, cells annotated as monocytes, dendritic cells, lymphoid, erythroid, or early progenitor populations showed no enrichment across the five datasets (**Fig. S5E**). We then subsetted the projected cells per study to assess cell state differences between cells variably harboring anellovirus transcripts (**Methods**). Differential expression analyses in the two datasets with the greatest number of anellovirus-positive cells showed an enrichment of mitotic cell cycle-associated genes, including *MKI67*, *RRM2*, *CENPE*, *KIF14*, and *CDC14B* (over-representation analysis *p* = 4.2×10^-20^ and *p* = 0.0032; **Fig. S5F**). As anelloviruses encode no polymerase and replicate using host machinery, we reasoned that the association likely reflected variable stages of the cell cycle. Indeed, annotating the Seurat-derived cell cycle stage showed a significantly elevated S/G2M occupancy, consistent with productive infection^56,57^ (**Fig. 5F**). Namely, 75% of anellovirus super-expressors (10+ UMIs) were in G2M, compared with 29% of cells carrying 1 to 9 UMIs or 19% of cells carrying none (beta regression; *p* = 8.5 × 10^-8^ and *p* = 0.00012; **Fig. 5F**). These findings suggest that proliferating cells may provide a permissive environment for anellovirus amplification, potentially linking viral activity to the availability of host replication machinery. Together, these analyses identify granulocyte–monocyte progenitors as a prominent site of anellovirus transcription in the sampled bone marrow datasets, consistent with prior detection of anellovirus DNA in granulocytes^4^.

### *Alphatorquevirus* ORF2 attenuates TNF-α response in myeloid cells

Our annotation of a potential anellovirus reservoir in myeloid progenitor cells motivated our focus on TNF-α and NF-κB signaling. Encouraged by recent work conducting viral ORF screens in human cells^58–61^, we sought to couple viral ORF perturbations with scRNA-seq profiling via Perturb-seq to discern the phenotypic effects of anellovirus proteins in single cells. As recent work has established ORF1 as the viral capsid^62^ and implicated ORF2 as a putative immunomodulatory agent^63,64^, we sought to characterize their functional perturbations in hematopoietic cell-line models. We selected ORF1 and ORF2 from high-and low-abundance strains across the three major viral genera and established a system to screen the transcriptional impacts of these proteins in single cells in response to cytokine stimulation (**Fig. 6A; Fig. S6A,B; Table S5; Methods**). We constructed a lentiviral vector with dual tetracycline-inducible transcriptional control and destabilization-domain regulation of protein stability, enabling precise control of anellovirus protein expression in human cells (**Methods**). We then transduced U937 and HL-60, common myeloid cell lines, and Jurkat, a common T cell line, with pools of anellovirus ORFs and monomeric enhanced GFP (meGFP) as a negative control, which we confirmed by flow cytometry and western blot analyses (**Fig. S6C-F**). Transduced cells were subsequently treated with TNF-α and/or IFN-γ, two cytokines upstream of pathways implicated in our biobank analyses, and profiled via scRNA-seq with custom probes to detect viral ORFs (**Fig. 6A; Methods**).

**Figure 6.**
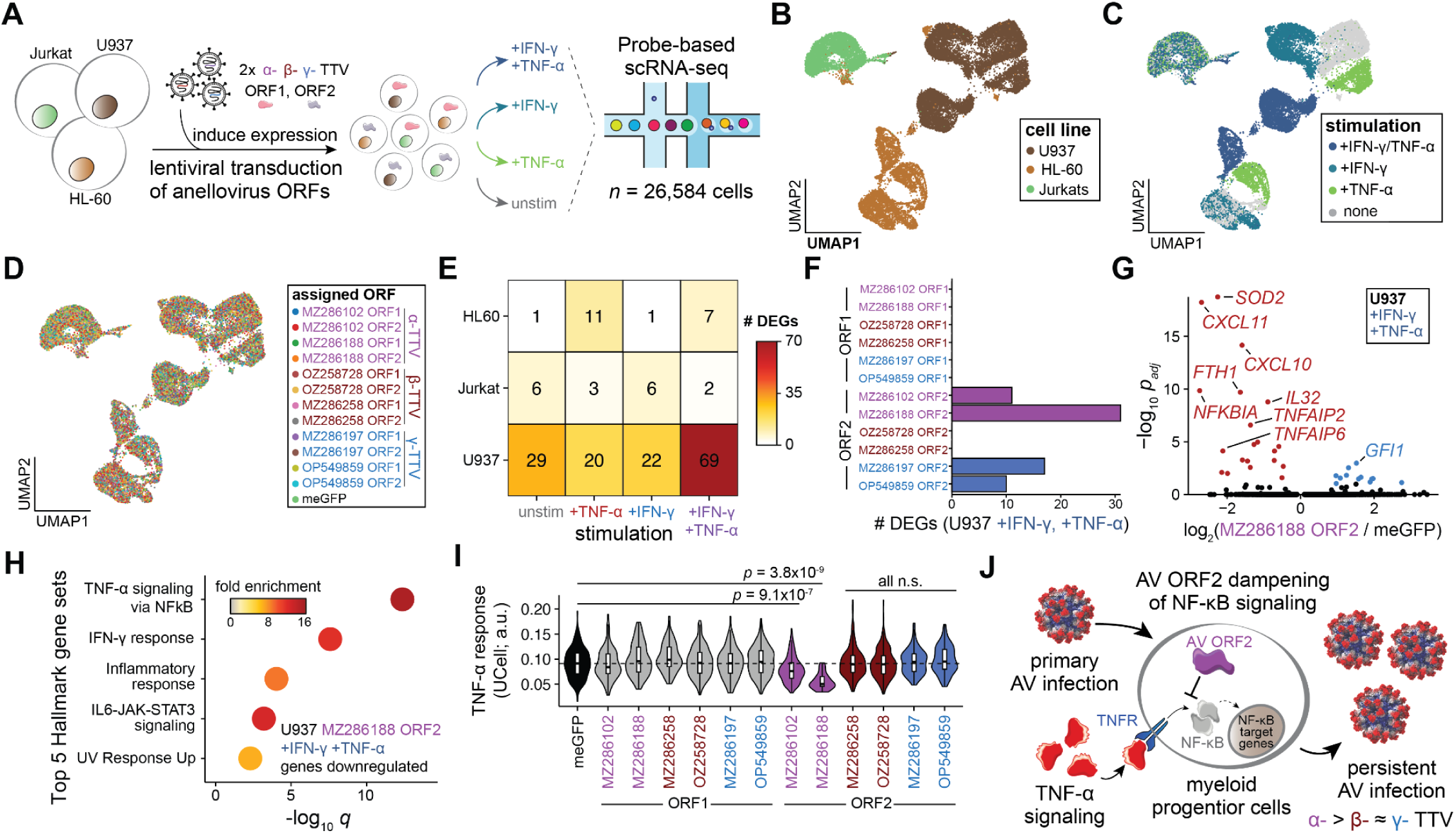
Characterization of anellovirus proteins via single-cell perturbations. A, Schematic of experimental design. Three cell lines were transduced with a pool of anellovirus ORF proteins, then variably stimulated before single-cell genomics profiling. **B,** UMAP embedding of single-cell data, colored by cell line. **C,** UMAP embedding colored by stimulation. **D,** UMAP colored by assigned ORF. **E,** Summary of the number of differentially expressed genes between anellovirus ORF proteins relative to the GFP control. **F,** Number of host genes differentially expressed following anellovirus ORF expression after IFN-γ/TNF-α in U937s. **G,** Differential gene expression summary of U937s stimulated with cytokines. Top associated genes are annotated. H, Gene over-representation analysis showing top 5 Hallmark gene sets with enrichments for cytokine stimulation in U937. **I,** Summary of TNF-α response in U937s treated with TNF-α. **J,** Schematic of hematopoietic populations regulating anellovirus persistence.

The two myeloid cell lines, U937 and HL-60, responded robustly to stimulation with TNF-α and IFN-γ, whereas Jurkat responded minimally (**Fig. 6B,C; Fig. S6G**). All anellovirus ORF or the meGFP negative control were readily detectable in all cell types, allowing for systematic differential expression analyses (**Fig. 6D; Methods**). Comparing each ORF to meGFP within a cell line and stimulation context, U937 cells expressing variable anellovirus ORFs showed the most differentially expressed genes (DEGs), and nearly all were attributed to the effects of our selected ORF2 proteins from the *Alphatorquevirus* and *Gammatorquevirus* genera (**Fig. 6E,F**). Notably, expression of *Alphatorquevirus* ORF2 significantly attenuated the expression of canonical NF-κB-inducible genes including *SOD2*, *NFKBIA*, *TNFAIP2*, *TNFAIP6*, as well as IFN-γ-inducible genes such as *CXCL10*, *CXCL11*, and *IL32*, supporting the suspected role of ORF2 as an intracellular immunomodulatory agent^63^ and implicating genes previously identified from our genetic association analysis (**Fig. 2**; **Fig. 6G, Fig. S6H**). Indeed, the top enriched hallmark gene sets included TNF-α signaling via NF-κB and IFN-γ responses (**Fig. 6H**), reflecting a convergent immunomodulatory axis from our GWAS and the effects of the *Alphatorquevirus* ORF2 protein. This effect of transcriptional dampening of the NF-κB response by TNF-α was apparent only for both *Alphatorquevirus* ORF2s proteins, but not for ORF2 proteins from the other genera or for any screened ORF1 protein (**Fig. 6I; Fig. S6I**). Instead, the effects of the *Gammatorquevirus* ORF2 proteins were distinct, as downregulated genes included promyelocytic/early granulocyte cell state markers *ELANE* and *CTSG* (**Fig. S6H**). Collectively, our data support a model in which anellovirus infection and persistence in myeloid progenitors are controlled by alterations in TNF-α-mediated NF-κB signaling, and these alterations may arise from host genetic differences or from modulation by anellovirus ORF2 (**Fig. 6J**).

## DISCUSSION

Emerging efforts have sought to map the abundance and diversity of the human virome along with the host factors regulating the persistence of trillions of infectious particles that cohabit the human body^22,24,25,37^. Because anelloviruses are the most abundant and diverse eukaryotic viral species in the virome^65^, mechanisms governing their infection and persistence over decades in the human body may generalize to other infectious agents that cause no overt phenotype. Here, we overcome the challenge of studying anelloviruses by accounting for their immense genetic diversity via sequence masking and pseudoalignment to produce a sensitive and specific viral pangenome suitable for retrospective quantification at scale. This approach enabled us to interrogate over 1.7 million sequencing libraries from primary human samples, spanning blood-based whole-genome sequencing (>1.1M; **Fig. 1**), targeted panel sequencing (>200,000; **Fig. 4d**), and single-cell RNA sequencing (>400,000; **Fig. 5**). Notably, our insights were derived from scalable reanalysis of libraries that were processed without specific intent to study anelloviruses or any other infectious agent, demonstrating the power of our approach to uncover novel biology from ordinarily discarded data.

While nearly every human has been infected by anelloviruses, our characterization shows that ∼16% of individuals in genetic biobanks have detectable viral DNA from discarded WGS reads. Consistent with anellovirus loads fluctuating with immune states, we estimate that ∼1-2% of donors in these biobanks exhibited active viral replication, as evidenced by high levels of anellovirus transcriptional activity based on RNA-seq. However, for most individuals, we interpret that anellovirus DNA detection in WGS data reflects a molecular phenotype of a subset of individuals with high levels of circulating anellovirus DNA when blood was drawn for WGS. Genetic association analyses of this variable anellovirus detection identified more than 120 loci genome-wide, over ten times the number reported previously^25^, revealing extensive host genetic control of viral persistence^25^. Genes near these implicated loci converge on immune-associated pathways of lymphocyte regulation and TNF-α-mediated NF-κB signaling, reflecting the altered immune surveillance that underlies elevated viral persistence in the population. Complementary analyses spanning genetic association analysis (**Fig. 2D**), plasma proteomics (**Fig. 3F**), and viral ORF screens (**Fig. 6H**) all converge on this predominant axis of TNF-α-mediated signaling, and we further tie attenuation of this axis to the functional activity of the anellovirus ORF2 protein.

Our annotation of donors in biobanks with detectable anellovirus DNA provides a framework for interpreting recent clinical associations of persistent anellovirus DNA detection. While our PheWAS implicates chronic conditions spanning cancer, kidney disease, and susceptibility to other infectious agents, we reason that anellovirus detection likely reflects an altered immune environment that permits expansion of an otherwise controlled member of the commensal virome. Plasma proteomic profiles suggest that individuals with persistent anellovirus DNA exhibit a peripheral signature consistent with altered immune regulation, including features of T cell dysfunction (**Fig. 3E**). We emphasize that these associations do not establish whether viral activity contributes directly to pathogenesis or results from disease and its treatment. Notably, while our human anellovirus pangenome substantially increased sensitivity for detecting viral persistence, individual species were observed too infrequently to adequately power strain-specific genetic or phenotypic association analyses. Instead, our data suggest that composite anellovirus burded may have prognostic value as an indicator of immune competence, and future work may enable strain-level association studies to refine specific disease associations. This outlook motivates the continued investigation of the blood virome as a component of high-dimensional profiling to improve risk stratification of complex disease^66^, including independent prospective evaluation.

Finally, through systematic reanalysis of all public, human scRNA-seq data in Logan^49^, we identified myeloid progenitor cells in the bone marrow as the most prominent site of anellovirus transcription (**Fig. 5**). While past work has implicated granulocytes as a major hematopoietic site of anellovirus replication^4^, other studies have also suggested T cells as a replication-competent reservoir^11,47^. While we cannot exclude the possibility that specific anellovirus strains infect and replicate in T cells, or that T cells serve as a reservoir for passive viral persistence, our data strongly support myeloid cells as a primary cell type for anellovirus replication in the bone marrow. Further, host transcriptomic analyses suggest that proliferating cells may provide a permissive environment for anellovirus activity. Together with our viral ORF Perturb-seq (**Fig. 6**), these observations support a model in which anelloviruses exploit permissive cellular states and attenuate immune signaling to promote persistence. These features are also relevant to emerging efforts to develop commensal virus-inspired gene-delivery vectors^21^ as the host determinants of persistence, cellular contexts of viral transcription, and immunomodulatory functions of viral proteins will all influence vector activity and host responses. Continuing to extend our computational and experimental approach across a broader array of species in the human virome will further reveal how cellular reservoirs and immune interactions determine whether viruses are cleared, persist, or expand in human health and disease.

## Acknowledgements

We thank members of the Norman, Dey, and Lareau labs for their helpful feedback, J. Cabral for helpful discussions, and B. Nalbant for help with flow cytometry. We acknowledge computing assistance from the Memorial Sloan Kettering High Performance Computing group. This work was supported by the Human Virome Program (U01AT012984 to C.A.L. and R.S.D.), P30CA008748 (J.C.G., T.M.N., K.K.D., and C.A.L.), a National Academy of Medicine Catalyst award (C.A.L.), and a Michelson Prize Next-Generation Grant (C.A.L.). R.S.D. and C.A.L. are supported by the American Brain Foundation’s Cure One, Cure Many award in Neuroinflammation. C.A.L. is further supported by R00HG012579, RM1HG011014, and R37CA303960. K.K.D. is supported by R00HG012203 and R01HG014008. T.M.N. is supported by R01HG014186. A.J.V. and C.A.L. are supported by a Geoffrey Beene Award. R.S.D. is supported by NIH DP5 OD036131, a Longevity Impetus Grant from Norn Group, Hevolution Foundation and Rosenkranz Foundation, and the Texas Children’s Hospital Research Vision Scholar program. J.C.G. is supported by T32CA254875. C.C.H. was supported by a Medical Scientist Training Program grant from the National Institute of General Medical Sciences of the National Institutes of Health under award number T32GM152349 to the Weill Cornell/Rockefeller/Sloan Kettering Tri-Institutional MD-PhD Program. E.M. is supported by F31HG014410. K.H. is supported by a Mildred Scheel Postdoctoral Fellowship (Deutsche Krebshilfe, 57856973). L.C.K. is supported by a Berlin Institute of Health MD Fellowship. S.S.N. and A.A.V. are supported by NSF Graduate Research Fellowships. R.C.A. is a Robert Black Fellow of the Damon Runyon Cancer Research Foundation (DRG-2462-22).

## Author Contributions

J.C.G., T.A.A., C.C.H., K.K.D., and C.A.L. conceived and designed the research. J.C.G., T.A.A., J.H.N., K.H., and C.A.L. led bioinformatics analyses with input from S.S.N., A.L., and A.A.V. C.C.H. led experimental work with input from A.J.V., L.C.K., R.C.A., E.M. J.C.U., P.P.F., T.M.N., and K.K.D. supervised aspects of the work. J.C.G., T.A.A., C.C.H., and C.A.L. wrote the manuscript with input from all other authors.

## Supplementary Tables

**Table S1. Summary of anellovirus detection genome-wide association studies and cohorts.**

**Table S2. Differential Olink proteomic signatures from anellovirus detection.**

**Table S3. Summary of PheWAS associations with anellovirus detection.**

**Table S4. Overview of SRA scRNA-seq accessions with detected anellovirus RNA.**

**Table S5. Details of anellovirus Perturb-seq experiment.**

## METHODS

### Description of genetic biobanks

Throughout this work, we analyzed five distinct repositories of whole-genome sequencing (WGS) data and others of bulk-or single-cell RNA sequencing. For each WGS repository, we aligned all genomes to the hg38 reference and accessed them through the respective analysis portal. Thus, we extracted unmapped reads using either the ‘GATK PrintReads-L unmapped’ or ‘samtools view-f 4’ commands from the hg38-aligned.bam/.cram files and created intermediate unmapped paired-end sequencing reads in.fastq.gz format. These paired reads were processed using the anellovirus references noted above, with kallisto for efficient read mapping. Subsequent downstream analyses were performed in the respective analytical environment hosting the individual-level genetic data, and meta-analyses were performed on variant-or summarized bucket-level analyses as we previously described for EBV.^22^

Our first biobank was the UK Biobank (UKB), a collection of nearly 500,000 individuals aged 40-69 years at the time of donor recruitment. Participants were enrolled in the UKB in the United Kingdom between 2006 and 2010 and were followed over time with updated information from self-reported surveys, linkage to death and cancer registries, and the collection of heterogeneous biomarkers, imaging data, and other phenotypes. Data were accessed using the UKB RAP under application numbers 16549 and 26041. Because the UKB is the best-annotated biobank, we performed several exploratory analyses before validating them in later cohorts. In particular, we assessed immune status at the assessment visit at which the blood sample was used for WGS. We considered five definitions separately; they were not mutually exclusive. We defined lymphopenia (n = 11,805) and neutropenia (*n* = 327) from the full blood count as a lymphocyte count below 1 x 10⁹ cells/L (UK Biobank field 30120) and a neutrophil count below 0.5 × 10⁹ cells/L (field 30140). Immunosuppressant use (n = 10,171) was defined from the regular medications recorded at the nurse-led verbal interview at the same visit (field 20003), counting systemic immunosuppressive agents: calcineurin inhibitors (ciclosporin, tacrolimus), antiproliferative agents (azathioprine, mycophenolate, methotrexate), mTOR inhibitors (sirolimus, everolimus) and oral glucocorticoids. Immunodeficiency (*n*=1,394) was defined by an ICD-10 immunodeficiency code (D80–D84) recorded at any time in linked hospital inpatient records. Transplantation (*n*=414) was defined from hospital procedure records (OPCS-4) of solid-organ transplantation (*n*=294), allogeneic hematopoietic stem-cell transplantation (*n* = 38), or autologous hematopoietic stem-cell transplantation (*n*=84), counted only if the procedure date was on or before the date of the sequenced blood sample; some participants had more than one transplant type.

Second, we used the All of Us (AoU) release v9, a collection of over 400,000 donors profiled with WGS. All of Us participants were recruited through a network of partners across the United States beginning in 2018. Data were accessed through the Researcher Workbench, approved by the All of Us Institutional Review Board (IRB #172034). All analyses in AoU replicated the UKB analyses, including correlates of phenotypic associations, Olink proteomic data analyses, and clinical trait associations, as well as the full genetic association studies for a meta-analysis.

Our third cohort was the Alliance for Genomic Discovery^67^. Participants were drawn from BioVU, Vanderbilt University Medical Center’s biobank of residual blood collected during routine clinical care in Nashville, TN, linked to de-identified, longitudinal EHR data rather than dedicated assessment-center visits. Data were accessed via the Illumina Connected Analytics platform.

As a fourth cohort for validation and multi-omic analyses, we reanalyzed the Genotype-Tissue Expression (GTEx) v10 cohort. For both WGS and RNA-seq from peripheral blood, we obtained mapped alignment files after obtaining dbGaP permissions for nearly 1,000 donors. Processed alignment files were fetched from the relevant Google Cloud Project bucket and further analyzed locally.

A fifth cohort, used as a silver-standard negative control for anellovirus detection, comprised 2,504 whole genomes profiled via the 1000 Genomes Project (1000G), collected from consenting donors worldwide. Because B cells derived from peripheral blood were immortalized into lymphoblastic cell lines (LCLs), detection of persistent anellovirus should be largely null. Thus, this cohort represented a highly diverse set of sequences that could be leveraged to assess the specificity of anellovirus assignment (i.e., at most, only minimal anelloviruses should be assigned to these unmapped reads). Aligned.cram files for the 1000G short-read WGS were downloaded from the International Genome Sample Resource and analyzed locally.

A sixth cohort to validate the association between anelloviruses and lymphoid malignancies was the Memorial Sloan Kettering (MSK) Cancer Center IMPACT cohort, which received ethical approval (protocol 12-245) under the Institutional Review Board (IRB), and all patients provided written informed consent for the use of their genomic data for research purposes^45^. We processed aligned sequencing reads from both tumor and matched blood-derived panel sequencing on the MSK Iris cluster. This included 205,156 samples from 87,166 patients, spanning 117,635 unique tumor sequencing libraries and 87,521 normal control samples (peripheral blood, saliva, nail clippings), with longitudinal information on survival, treatment status, and collection date.

### Human anellovirus reference assembly and characterization

We sought to develop a resource to classify human anellovirus reads that would serve multiple objectives. First, an ideal reference would include the breadth of genetic diversity observed in human anelloviruses. Second, the reference should contain minimal low-quality features that would lead to errant classifications of DNA reads as anellovirus. Third, the reference would contain only anellovirus sequences to ensure efficient runtime. Finally, the reference would be ideally indexed using k-mers for efficient data processing at the petabase scale. We verified that our human anellovirus pangenome achieves all four metrics.

To maximize anellovirus breadth, we queried NCBIVirus^68^ for all anellovirus species (TaxID = 687329) with an annotated human host and complete genome assembly, which extracted 3,545 unique strains. We then applied RepeatMasker v4.1.9 with human repeat sequence reference (-species 9606) to specifically mask any misassemblies and retain informative viral repeats. We reduced sequence redundancy using Vclust^30^, yielding 2,023 reference strains. This set met our first objective of capturing broad anellovirus diversity and was substantially larger than references used in previous anellovirus GWAS. To annotate the genera of all sequences, we extracted the predicted ORF1 coding region using EMBOSS^69^ with parameters for circular genomes and a minimum ORF size of one kilobase. Using these ORF1 sequences, we performed a multiple sequence alignment and clustered the trimmed sequences into a phylogeny via iqtree2^70^. We classified individual strains lacking an anellovirus genus by hierarchical clustering to the mode of their group, yielding the final annotations in **Fig. 1c**.

To minimize errant assignments, we hard-masked low-complexity sequences with dustmasker^31^. Because many anellovirus assemblies had low-complexity sequences, including AT-rich segments and contig ends, dustmasker hard-masked 4.7% of the original sequences. To verify that the masking removed errant read associations, we processed all unmapped reads from the 2,504 whole genomes from 1000G. We expected little or no anellovirus DNA in these samples, while their genetic diversity provided an opportunity to identify human sequences prone to misclassification. **Fig. S1E** shows that 7,493 reads were likely errantly assigned as anellovirus in our unmasked set of 2,023 reference strains, whereas only 2 reads were assigned after masking. We used NCBI BLASTn to assess these two reads; both returned individual anellovirus strains as the top match (**Fig. S1I**), suggesting that these 2 molecules, out of billions, may indeed be derived from an anellovirus.

The third criterion was to ensure that our reference could be used on its own. As our read assignment used only unmapped reads from an initial hg38 alignment, we reasoned that the vast majority of human-derived DNA had already been accounted for from the upstream alignment. To verify this, we reprocessed the full 1,011 GTEx genomes with the masked reference alone and an appended T2T reference to determine if any assigned reads would be ambiguous between human and viral. Our analysis showed only 1% of molecules were ambiguous, which we determined were largely discordant read pairs and likely reflected low-level artifacts, including PCR chimeras. We further confirmed the specificity of our k-mer assignments by computing the minimum Hamming distance between each 31-mer and its closest sequence in T2T, yielding a median of 6 and confirming minimal ambiguity in falsely classifying human sequences (**Fig. S1G**).

Finally, the fourth criterion was that reads could be analyzed using a k-mer-based pseudo-alignment workflow (rather than strict read mapping) to minimize runtime. As the GTEx (sensitivity) and 1000G (specificity) cohorts were both conducted using kallisto^32^ indices of 31-mers and pseudoalignment, and validated by NCBI BLASTn,^71^ we determined that our overall workflow was supported for scalable downstream analyses, while recognizing that these controls cannot exclude all sources of contamination or mapping ambiguity.

Because of cloud computing costs and complexity, we analyzed the three large biobanks (UKB, AoU, AGD) using only the human anellovirus pangenome. To verify the sensitivity and specificity of our reference, we first analyzed the GTEx and 1000G cohorts to assess the quality of our anellovirus classification workflow. For these two resources, we extracted unmapped reads and reprocessed them using four candidate anellovirus references. Explicitly:

-The ‘This work; unmasked’ was composed of the 2,023 de-duplicated strains after Vclust processing. This reference was indexed using kallisto^32^ for 31-mers, resulting in 4,800,623 sequences. Our analyses suggest that this reference contains many low-quality k-mers, resulting in substantial false-positive read assignment (see **Fig. S1E**).

-The ‘This work; masked’ comprised the same 2,023 deduplicated strains but with ∼4.7% of sequences hard-masked by dustmasker, resulting in 4,410,583 unique 31-mers. We used this as the primary reference throughout this work because it has been shown to be specific (**Fig. S1E**) and sensitive (**Fig. S1F**) for anellovirus assignment.

-The Qiu^23^ reference comprised 38 unique human anellovirus strains used in a recent anellovirus persistence GWAS, originally derived from a prior study. Although it contained considerably fewer strains, several had low-quality sequences, resulting in substantial erroneous classification (**Fig. S1E,F**).

-Finally, the Kamitaki^25^ reference comprised only 5 strains, selected for quality and maximal genetic diversity via a multidimensional scaling analysis. As these selected strains were of high quality, no off-target assignments were detected (**Fig. S1E**). However, the selection of only 5 strains led to limited sensitivity to detect diverse anellovirus species (**Fig. S1F,J**), limiting downstream inferences, including the axis of genetic persistence of these viruses (**Fig. 2b,c**). Specifically, the Kamitaki study performed extensive genome-wide association analyses for these strains and identified only seven conditionally independent genome-wide significant loci, providing a comparator for the combined effects of reference breadth, phenotype definition, and cohort composition.

Finally, we note that a fifth study, Sasa^12^, used a conceptually similar approach to ours by including a broad set of anellovirus reference genomes pulled from NCBI Virus. However, their approach did not mask low-complexity repeats as well and required more reads to assign positivity. Based on our analyses of positive and negative control cohorts (**Fig. S1**) and sensitivity for GWAS associations (**Table S1**), we determined that requiring higher read abundance and not masking the reference would similarly yield suboptimal results for downstream analyses.

For all cohorts, the primary association phenotype was detection of anellovirus DNA in WGS libraries, coded as positive when at least one read was assigned to the masked pangenome and negative otherwise. We use this binary annotation for ease of interpretation and for computing summary statistics such as population prevalence and based on enhanced statistical power from potential GWAS traits (**Table S1**).

### Genome-wide association studies

Motivated by recent work elucidating a heritable component of EBV persistence using unmapped sequencing reads^22,24,25,37^, we sought to establish a similar framework for anellovirus detection. Because these prior studies used a binary outcome of detectable EBV DNA from WGS data, we reasoned that we could apply a similar threshold to study host genetic factors regulating anellovirus persistence. To confirm that binarizing at one read per strain would suffice here, we first screened a panel of candidate phenotypes constructed from the same kallisto quantification in European-ancestry donors from the UK Biobank. These candidates varied along three axes: the required read support, from at least one read supporting a contig up to at least ten; the genus composition, testing each of the three principal genera (Alpha-, beta-and gamma-torquevirus) alone, in each pairwise union and in their full union; and the measurement scale, adding the total number of assigned anellovirus reads and the number of distinct assigned contigs as quantitative alternatives. Each candidate phenotyped was analyzed via the complete REGENIE^72^ association pipeline, and candidates were compared directly on the number of conditionally independent genome-wide significant loci recovered (**Table S1**). Binarization at one or more reads across the full union of the three genera recovered the most independent loci and thus was used as our primary phenotype. Index variants were identified by iteratively selecting the most strongly associated variant and excluding all remaining variants within 1 Mb. Index variant pairs within 3 Mb were then evaluated for conditional independence, as previously described ^73^.

Participants were partitioned into genetic ancestry strata within each biobank and analyzed independently, yielding 13 cohort-by-ancestry strata: five in the UK Biobank (European, African, South Asian, East Asian and admixed), six in All of Us (European, African, Admixed American, East Asian, South Asian and Middle Eastern) and two in the AGD (European and African). Analyzed samples ranged from 1,342 to 449,240 participants (**Table S1**). All three biobanks followed the same two-step whole-genome regression design implemented in REGENIE^72^. Step 1 fit leave-one-chromosome-out ridge predictors per stratum on directly genotyped autosomal variants with minor allele frequency >= 0.01, missingness <= 0.05, and Hardy-Weinberg equilibrium P > 1 x 10^-10^ with leave-one-out cross-validation. Step 2 tested imputed dosages under reference-first allele coding, retaining variants with minor allele frequency >= 0.005, missingness <= 0.01, and imputation INFO > 0.7. We tested the binary phenotypes with a Firth-penalized likelihood using an approximate correction. All models were adjusted for sex, age, age^2^, a sex-by-age interaction, and the first ten within-ancestry genetic principal components.

We harmonized per-stratum summary statistics across biobanks before meta-analysis, resulting in 22,244,013 variants, and combined them via a fixed-effect, inverse-variance-weighted meta-analysis in METAL^74^, weighting each stratum by the inverse of its squared standard error. We performed two meta-analyses: a European-ancestry analysis combining the European strata of the UKB, AoU, and the AGD (**Fig. S2A**), and a pan-ancestry analysis combining all 13 strata (**Fig. 2a**). Variants with heterogeneity degrees of freedom ≤ 1, *I^2^* > 75 or Cochran’s Q *p* ≤ 0.1 were removed, retaining 20,067,164 variants pan-ancestry and 10,211,284 in the European meta-analysis. From these, conditionally independent loci were defined among genome-wide significant variants (P < 5×10⁻⁸) by iterative 1 Mb clumping. Index variants within chr6:24,500,000–34,000,000 (GRCh38) were collapsed to the single most significant variant and labeled as HLA. For remaining index pairs 1-3 Mb apart, independence was assessed by an approximate conditional chi-square test using *r*^2^ from 1000 Genomes European populations queried through LDlink as previously described. This procedure yielded 74 conditionally independent loci in the European meta-analysis and 121 in the pan-ancestry meta-analysis. We applied the same procedure to each candidate phenotype to derive the selection criterion described above (**Table S1**).

To quantify the aggregate contribution of common variation, narrow-sense SNP heritability was estimated with BOLT-REML^75^ (v2.5) in European ancestry participants, separately for directly genotyped variants within the MHC (chr6:24–35 Mb, minor allele frequency ≥ 0.01) and for all directly genotyped autosomal variants outside it, adjusting for sex, age, age² and the first ten genetic principal components. We applied the same partition to EBV detection from our previous phenotype and to a previously published anellovirus GWAS, and reproduced estimates with GCTA-GREML averaged over ten random subsamples of 20,000 participants. We compared estimates using a two-sided Z test. To identify biological processes enriched among the associated genes, we computed per-gene association Z scores using MAGMA^76^ from the European meta-analysis, using a 1000 Genomes European linkage disequilibrium reference, a 0 kb gene window, and excluding genes within the MHC. Ranked gene-level Z scores were tested against Reactome pathways by gene set enrichment analysis (GSEA) via the fgsea package (gene sets of 15-500 genes, 10,000 permutations) with Benjamini–Hochberg correction. To assess shared genetic architecture with other traits, we estimated genetic correlations between the European anellovirus meta-analysis and the traits implicated in our PheWAS via the Complex Traits Genetics Virtual Lab^77^ (CTG-VL) panel using cross-trait LD score regression^78^, with significance declared at |Z| > 3 for Bonferroni correction across the queried traits.

### Correlates of anellovirus persistence

Because active DNA virus infections produce RNA (either from acute infection or reactivation), we characterized anellovirus-derived RNA from several sources. First, to validate the sensitivity of our pipeline, we accessed bulk RNA-seq data from patients with Long COVID profiled at Mt. Sinai, comprising 1,392 libraries, and reprocessed them using a transcriptomic reference of 34 strains^11,39^. Reads from this biobank were accessed from Synapse (syn35874390) and quantified using the published pipeline and our kallisto quantification with the human anellovirus pangenome reference, which were broadly correlated but with a ∼40% increase in sensitivity using the human anellovirus pangenome (**Fig. S3A**). To validate our interpretation of increased strain diversity and transcriptional expression in an immunosuppressed population, we reprocessed 19 transplant recipients and 19 controls where peripheral blood was profiled with RNA-seq over a short time course following receipt of the SARS-CoV-2 vaccine^42^ (**Fig. 3d)**. For both datasets, RNA-seq data was reprocessed using kallisto and our human anellovirus pangenome, and a reads per million (RPM) estimate was computed from the total number of assigned anellovirus reads over the number of reads in the RNA-seq library.

For the strain-diversity comparison (**Fig. 3c**), we retained participants with at least 10 total anellovirus reads and defined a major strain as one with at least 5 assigned reads. The interpretation of the effects was robust across different cutoffs for total anellovirus reads and the minimum requirement to define a well-detected strain. We modeled the number of anellovirus strains detected per participant with negative binomial regression (glm.nb in MASS) to account for overdispersion in strain counts. Models were fit among anellovirus-positive participants with one model per immunosuppression indicator, adjusting for log-transformed total anellovirus reads, age, age^2^, sex, their interactions, and genetic principal components, consistent with our GWAS analyses. Effects are reported as incidence rate ratios (the fold change in expected strain count) with 95% Wald confidence intervals.

### Plasma proteomics analysis

We used normalized protein expression (NPX) data from Olink plasma proteomics assays provided by the All of Us study. After excluding protein assays and samples that did not pass AoU-defined quality control standards, all participants without matching anellovirus quantification in blood, all participants with predicted ancestry other than EUR, all participants with non-binary gender or undefined age, the analysis was based on 2,761 samples and 5,413 profiled proteins. We used limma 3.62.2 to model protein expression from anellovirus-positivity (at least one anellovirus read) or binned anellovirus read count (0, 1, 2, 3-5, 5-10, 10-50, or 50+ reads), and used age, sex at birth and population-genetic principal components 1-15 as covariates.

### Phenome-wide association studies

The exposure throughout was the same binary detection call used for the genetic association analyses, consistent with our prior efforts with EBV^22^. The UK Biobank served as the discovery cohort and for time-to-event models; it was restricted to 449,240 European-ancestry participants with an anellovirus positive-or-negative assignment, complete covariate data, and a sequencing date for the time-to-event analyses. Within this cohort, 33,663 participants (7.5%) carried detectable anellovirus DNA. Clinical outcomes were defined from ICD-10 codes drawn from six linked clinical sources: hospital inpatient main and secondary diagnoses (fields 41202 and 41270), primary and contributory causes of death (40001 and 40002), cancer-registry site and histology codes (40006, 40011 and 40012), and the harmonized first-occurrence fields (p130xxx). Self-reported illness (20001, 20002 and 6150) and derived data fields defined neither cases nor controls, so that every association rests on a clinically recorded diagnosis. Each ICD-10 entity was defined at three granularities: the 3-to 5-character code, the block, and the chapter, and tested both across all sources and within each source separately, because ascertainment differs between them. Traits with significant sex-bias were downsampled as previously described^43^. Cases were assigned by prefix matching, with chapters and blocks expanded to their constituent code ranges beforehand. Controls excluded any participant with a diagnosis elsewhere in the phenotype’s own ICD-10 chapter, and any carrying the phenotype itself under a permissive definition that additionally counts self-reported and derived sources. This yielded 12,661 binary phenotypes with at least 10 cases in which Firth regression converged to an estimate.

Each phenotype was regressed on anellovirus detection using Firth’s penalized likelihood logistic regression, implemented with logistf through the phewas_ext function of the PheWAS R package, adjusting for sex, age, age², a sex-by-age interaction, and the first ten genetic principal components, the same covariate set used in the genetic association analyses, and at a Bonferroni-adjusted threshold for significance, 58 tests were significant. Because the same clinical entity is represented at the code, block, and chapter level and in several ascertainment sources, we collapsed these 58 significant associations into 16 distinct traits, as summarized in **Table S3**. We repeated replication association analyses in AoU and AGD, on European-ancestry participants and using each biobank’s linked electronic health records, with the same phenotype logic, control definition, covariate set, and Firth model. Because the three cohorts code diagnoses at different granularities, the significant UK Biobank traits were matched to their AoU and AGD counterparts on ICD-10 identity alone at 3-character granularity. We considered a trait replicated when the matched counterpart showed a concordant effect direction and p < 0.05.

Because anelloviruses were unlikely to be a causal infection for the associated traits, we reasoned that the value of the associations may lie in prospective detection for patient stratification; hence, we emulated this with longitudinal association testing in the EUR UKB, where phecodes have been accumulating for over 15 years^79^. We excluded participants with a recorded diagnosis on or before the assessment-center visit at which the sequenced sample was drawn as prevalent cases, and followed the remainder to the first subsequent record of that diagnosis, to death, or to the end of follow-up in 2025. We tested anellovirus detection as a predictor of incident diagnosis using Cox proportional-hazards regression, adjusting for log-transformed total unmapped read count as a sequencing-depth covariate, age, sex, and the first ten genetic principal components. We show unadjusted event-free survival using the Kaplan-Meier method with survfit from the R ‘survival’ package in the same cohort, with the same prevalent-case exclusion. Curves are plotted with 95% confidence intervals and accompanied by numbers at risk. Because the Kaplan-Meier estimator is unadjusted, we present it as a descriptive companion to the Cox models, and we derive statistical comparisons between strata from the corresponding covariate-adjusted hazard ratio rather than a separate log-rank test.

### Correlates of anellovirus in MSK-IMPACT cancer sequencing data

Anellovirus reads were mapped from unmapped reads in MSK-IMPACT targeted sequencing data of tumor and normal samples as described above. Anellovirus-positivity was determined as at least one anellovirus read per sample. Samples were linked to clinical annotations using MSK-internal cBioPortal data, including patient sex, diagnosis, highest recorded stage, overall survival from the time of sequencing of the first sample, pretreatment information prior to MSK, treatment timelines, assay version, and sample sequencing coverage. Age at sequencing was not available due to data protection measures. Exact annotation of material and sampling times for normal samples was not consistently available; we therefore used the first sequenced tumor sample of each patient for clinical analysis. We used the OncoTree ontology to merge diagnostic groups with fewer than 50 patients (for clinical correlations) and 500 patients (for prevalence barplots). Pretreatment status was inferred from (a) pretreatment information prior to MSK and from (b) an annotated treatment start date of more than 7 days prior to specimen date. For survival analysis, we considered the first sequenced tumor sample to determine anellovirus status of each patient and analysed overall survival from the day of sequencing. We used multivariable Cox proportional-hazards regression, stratified by diagnosis and stage, and included anellovirus positivity, prior treatment, sex, assay version (MSK-IMPACT gene panel) and sequencing coverage as covariates.

### Identification of anellovirus-positive scRNA-seq data with Logan

Because anellovirus tropism has been reported in both T cells and myeloid populations, we sought to refine the populations that could support anellovirus infection using a broad range of publicly available scRNA-seq samples. To achieve this, we used the index of our human anellovirus pangenome and aligned contigs from 3.2 petabases of human single-cell sequencing assembled as part of the Logan v1.2 project.^49^ This was achieved by identifying single-cell RNA-seq libraries using the SRA metadata ‘librarysource == “TRANSCRIPTOMIC SINGLE CELL” & organism == “Homo sapiens”’, which contain anellovirus sequences for downstream reprocessing and single-cell viral quantification. Once these SRA accessions were identified, we aligned all Logan contigs (v1.2 release) against the human anellome reference using minimap2 v.2.30^80^ with the short-read preset (-x sr), with SAM output (-a) restricted to sequences with reported alignments (--sam-hit-only), and extracted individual contigs with sufficient alignment. Petabase-scale processing was performed on Amazon Web Services using AWS Batch and Docker containers executed on EC2 c5d-family Spot instances. We identified top-expressing samples by computing a Logan contig nomination score, defined as the total number of aligned bases multiplied by the ratio of aligned bases to total contig length. This value nominates long alignment tracks that represent a high percentage of the total contig, while still yielding positive values for shorter contigs with significant alignments. Because most top accessions were derived from bone marrow or blood (**Fig. 5b**), we estimated annotation enrichment using Fisher’s Exact Test, which showed clear enrichment of bone marrow RNA-seq libraries expressing anellovirus RNA (**Fig. S5A**).

We assessed the highest-expressing anellovirus-positive datasets by downloading the raw.fastq files from the SRA for all identified samples in the study. Using these sequencing reads, we ran two parallel workflows. First, to resolve single-cell anellovirus expression, we processed the.fastq files with kallisto|bustools, specifying the sequencing chemistry and using our human anellovirus pangenome with the hg38 genome as a ‘decoy’. Second, we used CellRanger v10 with the standard hg38 reference to reprocess all samples into human scRNA gene expression matrices. For bone marrow-derived populations, we projected the processed single-cell count data from the corresponding accessions onto a recently described harmonized MarrowMap atlas.^50^ For peripheral blood populations, we reclustered and annotated them with marker genes, as existing references lack granulocytes in the embedding, and we anticipated the highest anellovirus loads in neutrophil populations based on prior data. We imported the two matrices into Seurat v5^81^ with default settings to compute the cell cycle score, and used MarrowMap^50^ to annotate per-cell two-dimensional UMAP coordinates and predicted cell-type labels for downstream analyses. For downstream analyses, we observed that our results were robust to the choice of any individual cutoff value. To determine differential gene expression features between anellovirus super-expressors and uninfected cells, we subsetted to myeloid populations from the two datasets with the largest numbers of anellovirus-positive cells and used the FindMarkers function in Seurat. This yielded differentially expressed genes that included several cell cycle members (**Fig. S5F**). We computed the proportion of G2M cells within the myeloid compartment stratified by bins of anellovirus expression of exactly 0 viral UMIs, 1-9 UMIs, and 10+ for anellovirus super-expressors (**Fig. 5f**). Statistical inference was performed via a beta regression with the study label as a covariate.

### Anellovirus ORF expression and profiling

We constructed a lentiviral ORF expression vector to dual-control anellovirus ORF expression via doxycycline-inducible transcriptional activation and Shield1 ligand-inducible stabilization of N-terminally destabilization domain (DD)-tagged ORFs. Anellovirus ORF1s and ORF2s from each genus at the high and low ends of biobank-determined population prevalence were selected for inclusion in our single-cell perturbation experiment (**Fig. S6A**). For visualization purposes, ORF1 and ORF2 amino acid sequences for the six selected anellovirus strains were aligned separately using AlignSeqs from DECIPHER^82^ with default parameters (**Fig. S6B**). For each alignment column, the consensus residue was defined as the most frequent non-gap amino acid, and every residue was then classified as matching consensus, differing from consensus, or a gap. In total, this represented six ORF1s and six ORF2s, along with monomeric enhanced GFP (meGFP) as an internal negative control. ORFs were tagged with HA for detection of protein expression by Western blot (**Fig. S6F**). Tags were placed at the C-terminus of ORF2 and meGFP, and at the N-terminus of ORF1, as ORF1 is known to undergo C-terminal proteolysis in human cells^62^.

Constructs were ordered as synthetic DNA and cloned individually into the lentiviral ORF expression backbone using Gibson assembly. Lentivirus was produced in Lenti-X 293T cells, in two separate pools for ORF1 and ORF2, with meGFP spiked into each pool at 5% DNA by mass. Lentivirus-containing supernatant was collected at 48 and 72 hours post-transfection, pooled, and concentrated using a Lenti-X concentrator. ORF1 and ORF2 lentivirus were titered in U937, HL-60, and Jurkat cell lines, and transduced at a multiplicity of infection (MOI) of 0.3 (**Fig. 6a**). Cells were subsequently FACS-sorted for BFP+ (transduced) and recovered until sufficient cell numbers were attained for the downstream experiment (**Fig. S6C**). We designed two pairs of custom Flex probes per ORF using a custom probe design pipeline.^83^

One million cells from each transduced cell type, representing 6 cell pools total (ORF1 vs. ORF2, three cell lines), were induced for ORF expression with 1 μg/mL doxycycline and 1 μM Shield1 for 24 hours. We confirmed induction by flow cytometry and anti-HA Western blot at 24 hours. We then pooled ORF1-and ORF2-transduced cells from the same cell type and split them four ways for the following stimulations: unstimulated, TNF-α 20 ng/mL, IFN-γ 50 ng/mL, and combined TNF-α 20 ng/mL + IFN-γ 50 ng/mL. Cells were stimulated for 6 hours at 37°C, then fixed overnight at 4°C per the 10x GEM-X Flex Gene Expression protocol. The following day, cells were prepared for long-term storage and frozen at-80 °C. Upon thawing, cells from the same stimulation conditions were pooled 1:1:1 to a final total of 350,000 cells, as cell types could be deconvoluted in downstream analyses. We hybridized the 10x Flex v2 whole-transcriptome atlas (WTA) and custom viral ORF probes to the four multiplexed samples overnight at 42°C. The six strains were MZ286102 and MZ286188 (*Alphatorquevirus*), OZ258728 and MZ286258 (*Betatorquevirus*), and MZ286197 and OP549859 (*Gammatorquevirus*), each represented by ORF1 and ORF2.

We prepared a four-plex 10x Genomics Flex library according to the manufacturer’s instructions using the Flex Apex Probe set and sequenced it on the Element Aviti with standard dual indexing and demultiplexing. Raw.bcl files were processed using CellRanger v.10.0, and the resulting.fastq files were quantified for the human v2 probe set using default parameters for the CellRanger pipeline with the hg38 (human) reference. We performed downstream analyses, including cell filtering, marker-gene analyses, and visualization, using Seurat v5. In brief, we identified cells by passing the CellRanger knee call and meeting additional quality-control standards: at least 1,000 UMI counts, at least 500 genes detected, and no more than 5% mitochondrial RNA abundance (standard thresholds for scRNA-seq analyses). We considered cells assignable if exactly one ORF had at least four UMIs, or if the most abundant ORF had at least two UMIs and accounted for at least 75% of all perturbation-associated UMIs in the cell. Cell doublets were removed with either multiple high-confidence ORF detection assignments or based on heterotypic doublets using DoubletFinder^84^. We performed all differential expression and marker-gene analyses using the FindMarkers function in Seurat, comparing against the meGFP-assigned cells. TNF-α-response scores were computed with UCell^85^ using the set of genes in the Hallmark gene set.

## Lead contact and materials availability

The lead contact is responsible for supplying any and all materials related to this work upon reasonable request.

## Resource availability

Code to reproduce and use the human anellovirus pangenome is available on GitHub: https://github.com/clareaulab/anellovirus_reference.

## Code availability

Source code for the Logan cloud computation system is available at https://gitlab.pasteur.fr/rchikhi_pasteur/logan-analysis. Code to reproduce custom anellovirus Perturb-seq analyses is available at https://github.com/clareaulab/anello_perturbseq_v1.

## Data availability

Single-cell sequencing data generated as part of this work are available on the Gene Expression Omnibus at accession **GSE347221** with access token **sdmxqomehhwzvix**.

Genome-wide association study summary statistics can be downloaded from Locuszoom for both the European (https://my.locuszoom.org/gwas/307635/?token=f88c000df25647deae22c9fdbc7d3b06) and pan-ancestry (https://my.locuszoom.org/gwas/669893/?token=aee551c04a48482dbded3694d29e6675). Individual-level genetic and phenotypic data, including viral phenotypes, must be requested following an access agreement per each biobank.

---

**Figure S1.**
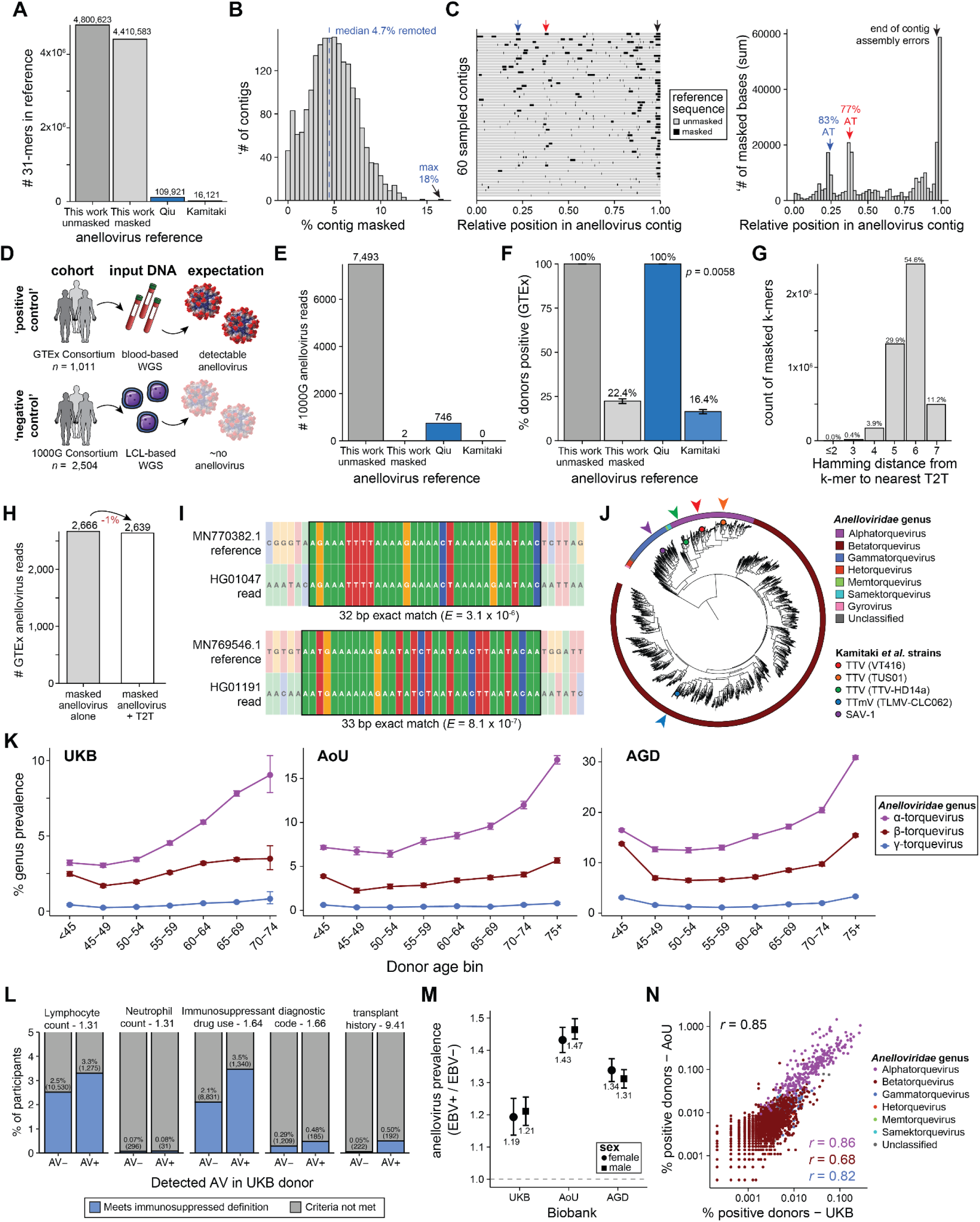
Supporting information for human anellovirus reference curation and biobank analyses. A, Quantification of anellovirus-associated 31-mers from indexing various viral references. Qiu^23^ and Kamitaki^25^ used 38 and 5 strains, respectively, to build their reference sets. **B,** Summary of impact of masking per contig, noting the median percentage of bases masked (4.7%). **C,** Visualization of loci masked, including two AT-rich regions noted in red and blue. **D,** Schematic of two cohorts, Genotype-Tissue Expression (GTEx) and 1000 Genomes Project (1000G), input material for WGS, and expected anellovirus detection. **E,** Quantification of anellovirus from 1000G (negative control), showing the number of reads quantified from the unmapped fraction. **F,** Percent positive donors from GTEx (positive control). Error bars: standard error of the mean; Statistical test: Fisher’s Exact test comparing the masked reference and Kamitaki reference. **G,** Summary of distinct anellovirus 31-mers to nearest match in the T2T reference genome. **H,** Number of anellovirus reads from the unmapped fraction using the masked human anellovirus pangenome with or without additional T2T inclusion in the reference. **I,** Characterization of 2 anellovirus-assigned reads from the 1000G via the masked human anellovirus pangenome. *E* values computed from NCBI BLASTn. **J**, Annotation of 5 strains from Kamitaki *et al.* onto the hierarchical clustering from Fig. 1c. **K,** Biobank summary of anellovirus detection stratified by donor and anellovirus genus. **L**, Immunosuppressed donor annotation stratified by anellovirus detection in the UKB for five definitions. The effect size (ratio of prevalences) is noted next to the title of each set. **M**, Ratio of prevalences of anellovirus detection comparing EBV-positive and negative donors stratified by sex and biobank. **N,** Corroboration of strain-level quantification between the AoU and UKB biobanks. Values show Spearman correlation overall (black) and for each major genus (colors).

**Figure S2.**
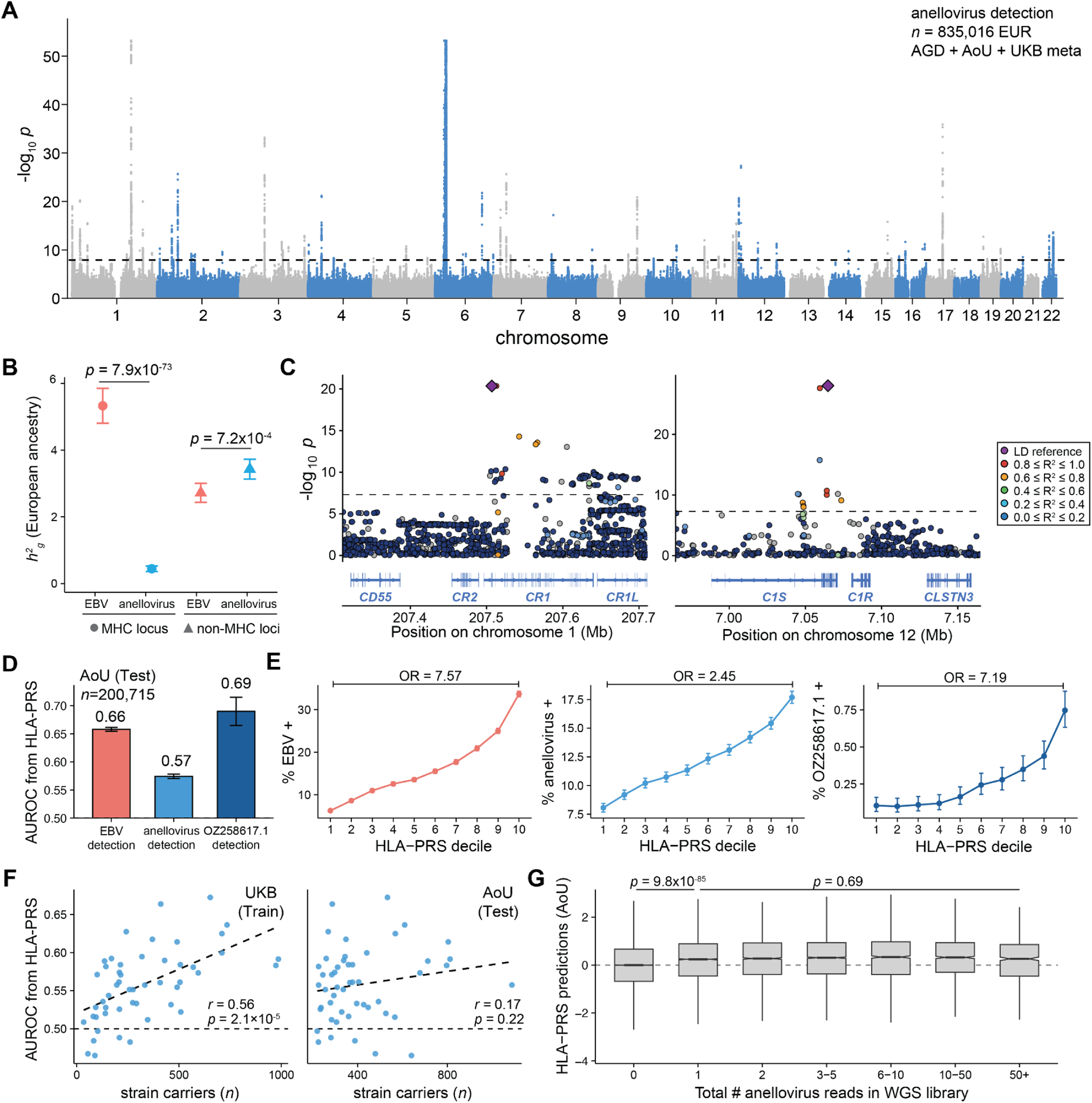
Supporting information for genetic association analyses. A, Manhattan plot of pan-European anellovirus DNA detection GWAS. **B,** Estimated heritability partitioned by inclusion or exclusion of the MHC locus for detection of EBV or anellovirus. Statistical test: two-sided Z test. **C,** Locus zoom plot of selected loci near the complement family genes. **D,** Performance of HLA polygenic risk score model for EBV detection, anellovirus detection, and a specific anellovirus strain, OZ258617, for the All of Us (AoU) cohort. **E,** Predicted deciles of HLA polygenic risk with proportion positive for each viral species. Odds ratios between the top and bottom deciles are shown. **F,** Performance of detection prediction among the 100 most abundant strains in training data (UKB; left) and test data (AoU; right). Statistical test: Pearson correlation. **G,** Performance of HLA-PRS predicting anellovirus detection. Statistical test: Mann-Whitney test.

**Figure S3.**
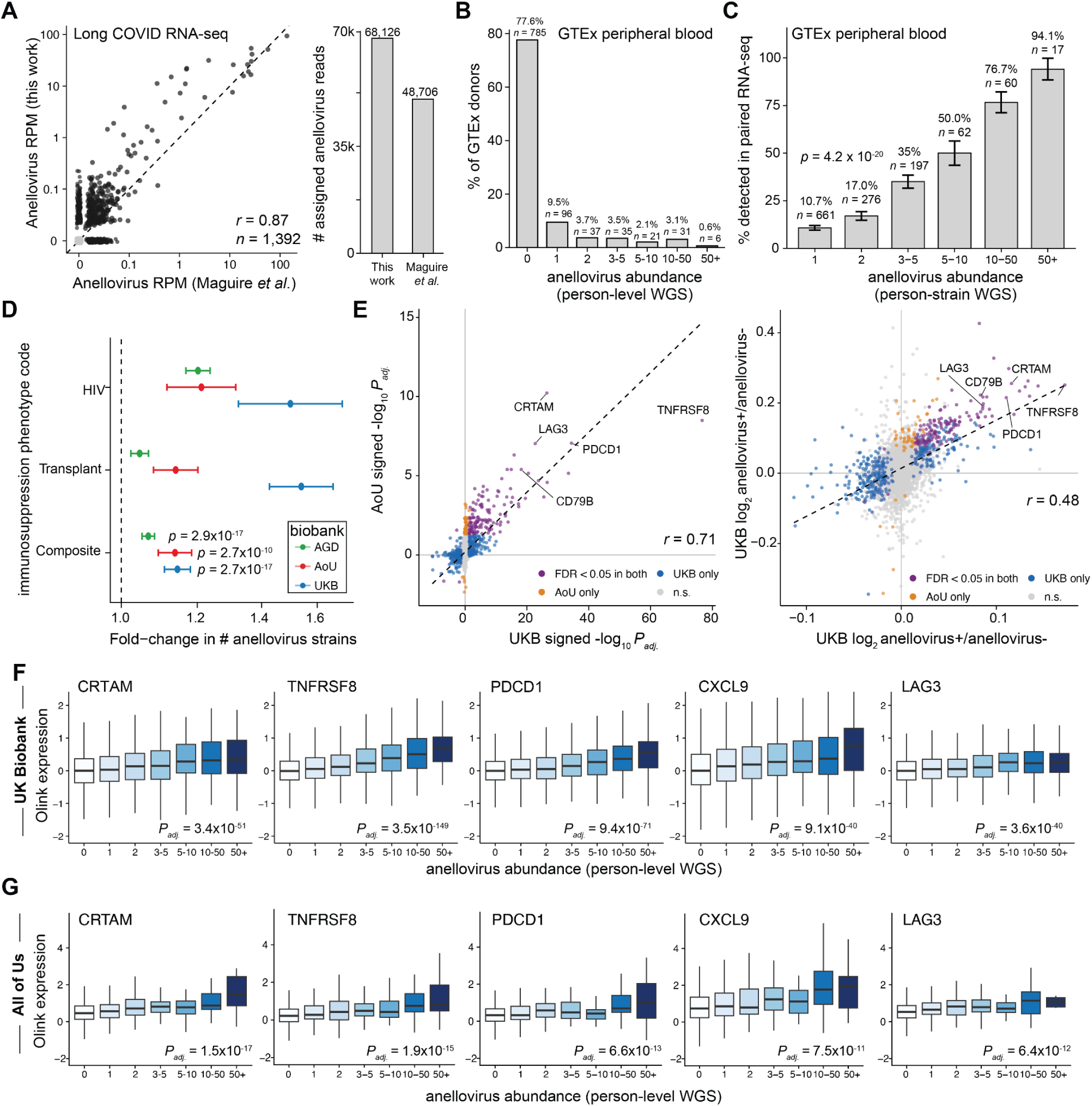
Supporting information for phenotypic correlations with anellovirus detection in large biobanks. A, Characterization of per-library anellovirus reads per million (RPM); data and comparator method from Maguire *et al.*^11^. **B,** Number of anellovirus reads per donor in the GTEx cohort using peripheral blood WGS. **C,** Summary of anellovirus transcriptional activity. Binned donor-strain WGS reads were tested for evidence of RNA in matched GTEx RNA-seq data. Statistical test: Cochran-Armitage trend test. **D,** Estimation of increased anellovirus strain abundances over variable immunosuppressive agents. Circles are point estimates from a generalized linear model; error bars reflect 95% confidence intervals. P values are from Wald statistics of a negative binomial regression. **E,** Concordance of Olink proteomic signatures between biobanks. Correlations are Spearman correlation statistics. **F,** Dose-response of top Olink differential proteins. Statistical test: limma linear model with Benjamini-Hochberg multiple testing correction. **G,** Same as **F** but for the All of Us Olink data (*n*=2,761 samples).

**Figure S4.**
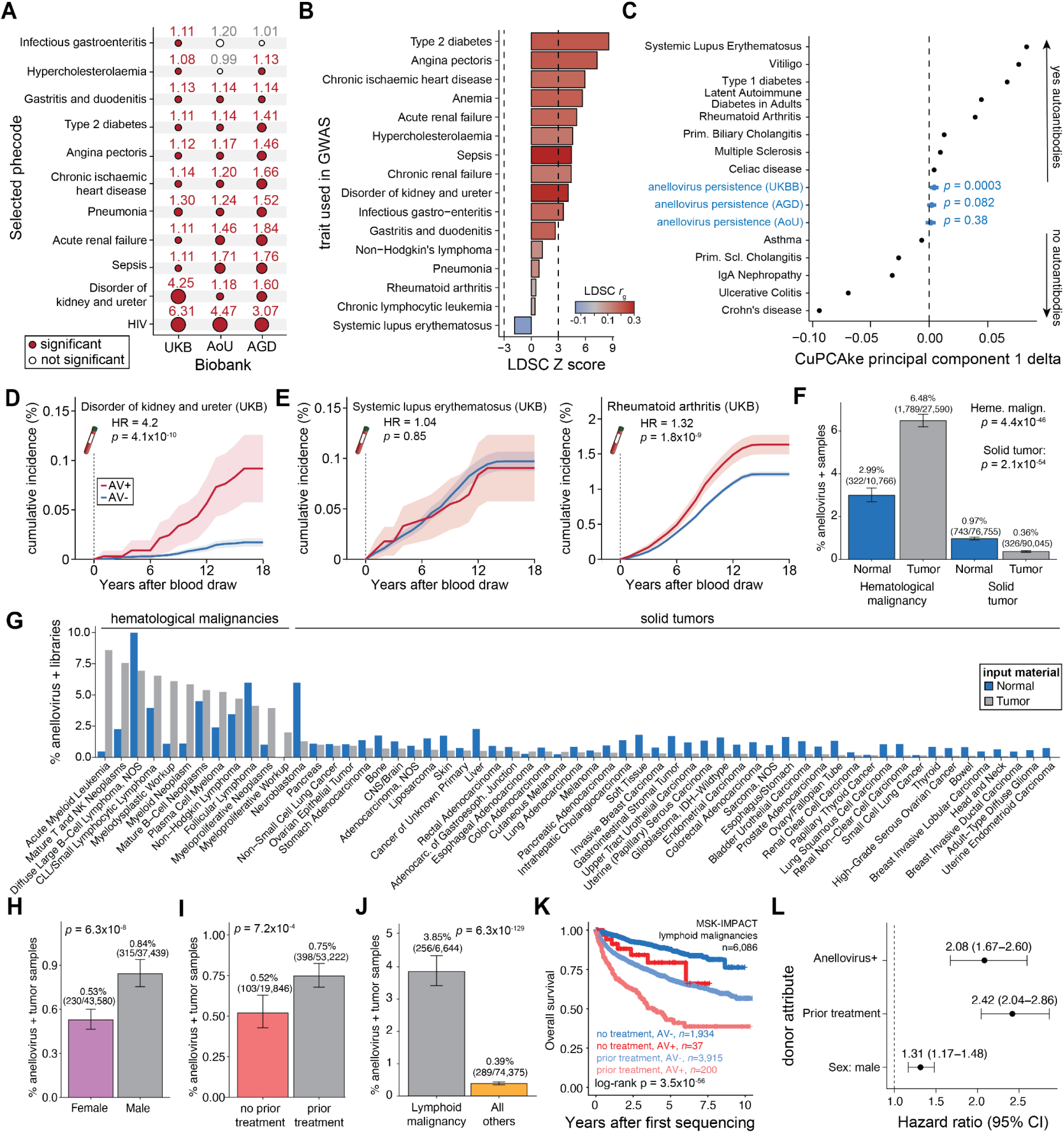
Supporting information for phenotypic associations. A, Comparison of effect sizes and significance for selected traits across the three genetic biobanks. Values represent odds ratios of association, and significance shows Bonferroni-adjusted associations. **B,** Genetic correlation of anellovirus persistence and PheWAS-significant traits. Dotted lines reflect Bonferroni-adjusted significance thresholds. **C,** Principal component analysis and projection of anellovirus persistence summary statistics on complex immune-mediated diseases via cupcake^44^. Statistical test: Z test of principal component projection score. **D,** Kaplan-Meier incidence curves for disorders of the kidney and ureter from EUR donors in the UKB. Statistical test: Cox-Proportional Hazards model. **E,** Same as **D** for systemic lupus erythematosus and rheumatoid arthritis. **F,** Percentage of anellovirus-positive MSK-IMPACT samples in hematological vs. other malignancies. Bars show Wilson 95% confidence intervals. Statistical test: two-sided Fisher’s exact tests. **G,** Percentage of anellovirus-positive tumor (grey) and normal (blue) samples by OncoTree diagnosis (diagnoses with <500 patients collapsed). **H–J**, Percentage of anellovirus-positivity by H, sex; **I,** treatment status prior to sequencing; and **J,** tumor type. Bars show Wilson 95% confidence intervals. Statistical test: two-sided Fisher’s exact tests. **K**, Kaplan-Meier analysis of overall survival from first sequencing for lymphoid malignancies, stratified jointly by anellovirus positivity and prior treatment status. P-value from global log-rank test across all groups. **L**, Forest plot of hazard ratios (95% confidence intervals) from a multivariable Cox proportional hazards model of overall survival in lymphoid malignancies.

**Figure S5.**
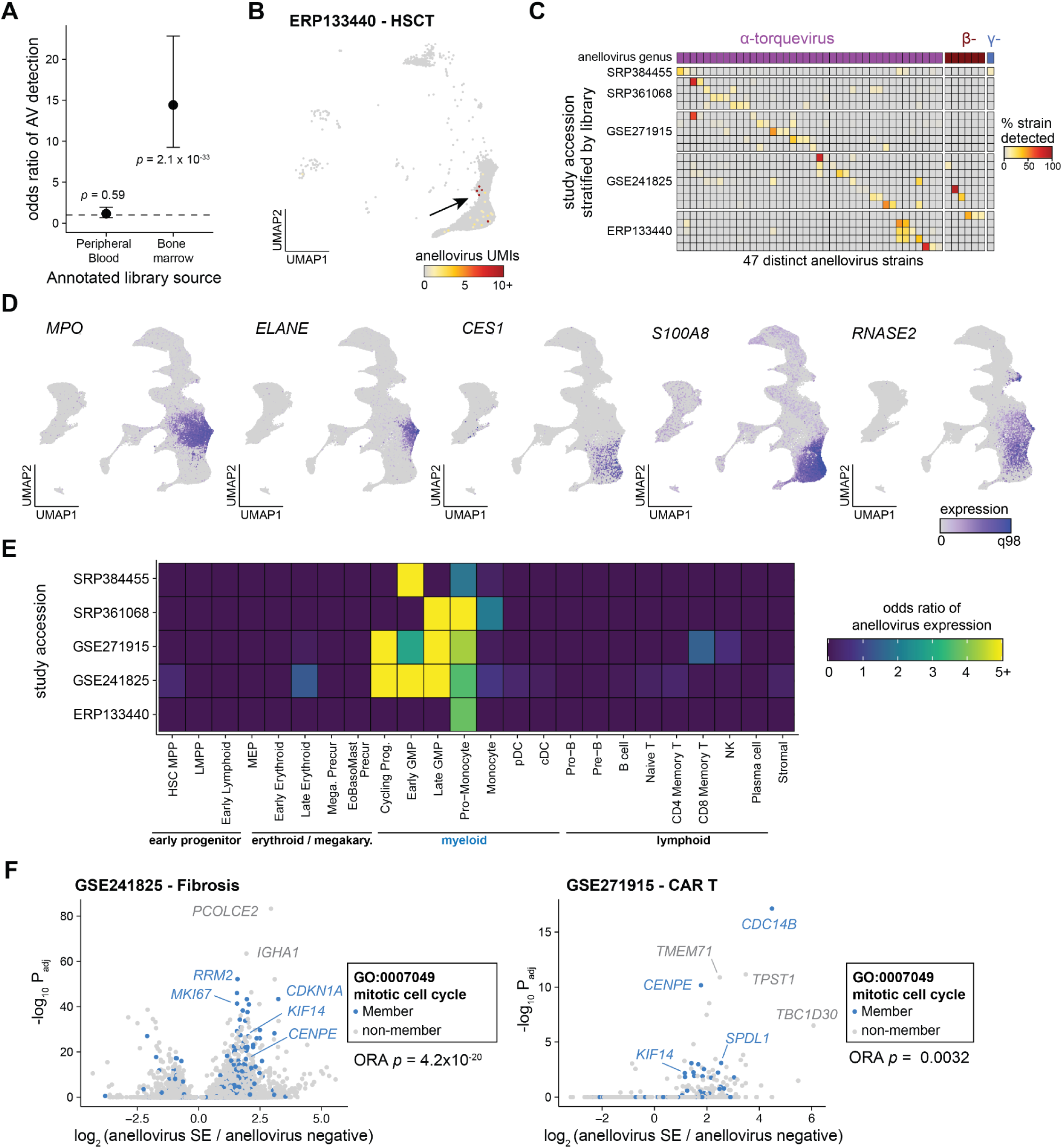
Supporting information for petabase-scale mapping of anellovirus detection in single-cell data. A, Relative enrichment of scRNA-seq tissues by anellovirus (AV) detection by tissue source from single-cell data mining. Statistical test: Fisher’s exact test. **B,** Projection of a hematopoietic stem cell transplant (HSCT) scRNA-seq dataset showing abundance of anellovirus reads in individual cells. **C,** Donor-and strain-level abundance across the Logan-identified contigs^49^. **D,** Notable marker genes in myelocytic precursor populations overlapping the anellovirus-positive cells. **E,** Study-and population-level enrichment (odds ratio) of anellovirus-positive cells. **F,** Differential gene expression of myeloid populations stratified by anellovirus super-expressors (SE) or negative cells, highlighting genes involved in the mitotic cell cycle. Statistical test for over-representation analysis (ORA): two-sided hypergeometric test of differentially expressed genes (*P*adj < 0.01).

**Figure S6.**
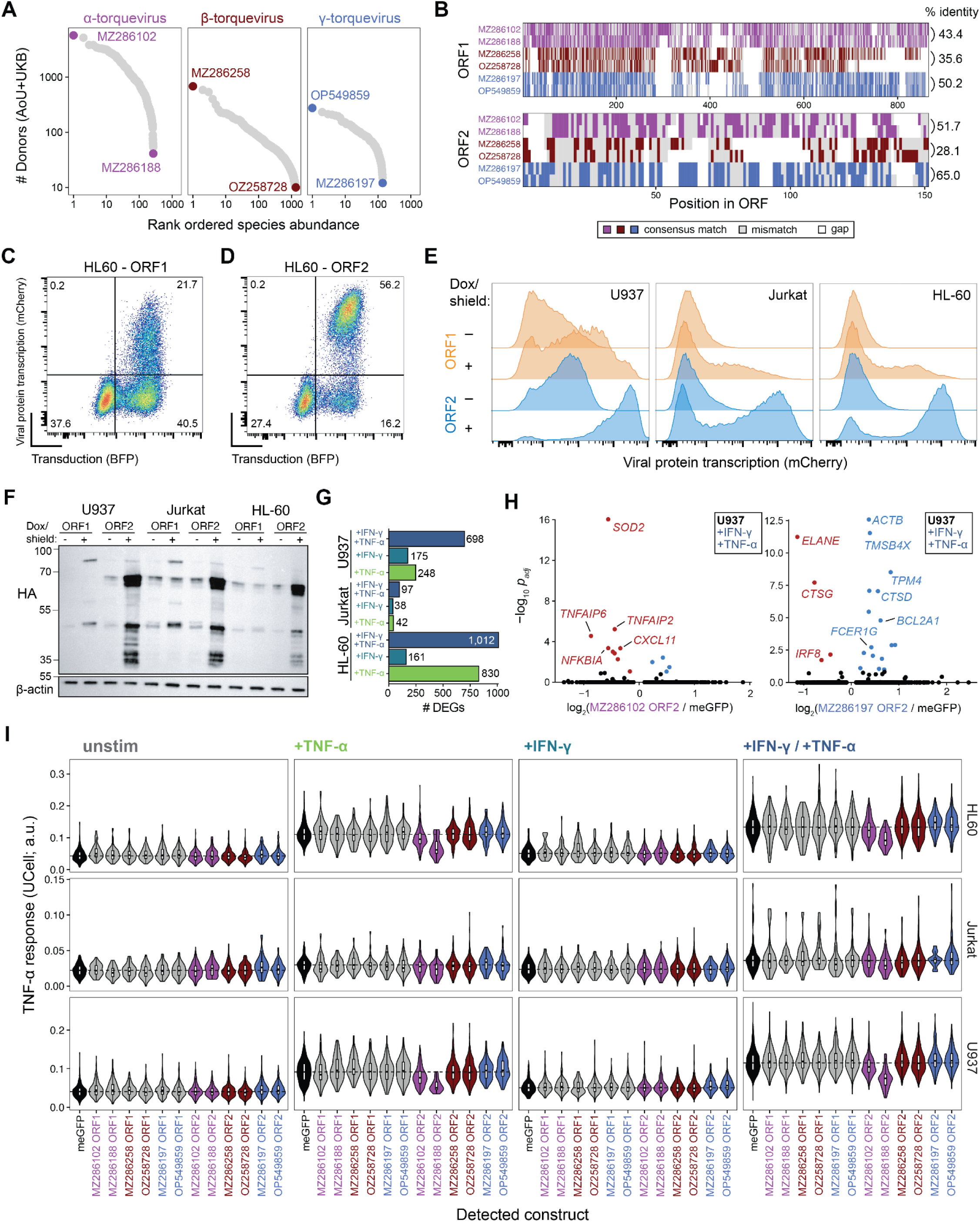
Supporting information for single-cell analyses. A, Selection of 6 strains for functional characterization. **B,** Multiple sequence alignment of ORF1 (top) and ORF2 (bottom) proteins, colored by genus. **C,** Representative flow cytometry plots marking transduction (BFP) and induction of viral protein transcription (mCherry) for ORF1. **D,** Representative flow cytometry for ORF2. **E,** mCherry induction before and after doxycycline/Shield1 treatment for each cell line and anellovirus proteins ORF1 and ORF2. **F,** Confirmation of HA-tagged anellovirus protein and meGFP expression after doxycycline/Shield1 treatment. **G,** Number of differentially expressed genes (*P*adj < 0.01; abs(log2FC) > 1) comparing each cell line; **H,** Volcano plots of ORF2 TTV strains. Left: MZ286102 ORF2 strain (α-TTV) compared to meGFP; right: MZ286197 ORF2 strain (γ-TTV) compared to meGFP. Statistical test: Mann-Whitney U-test with Benjamini-Hochberg multiple testing correction. **I,** TNF-α response module score across all cell lines, stimulations, and ORF combinations. Non-null effects are highlighted with arrows.

